# Macromolecular interactions dictate Polycomb-mediated epigenetic repression

**DOI:** 10.1101/2025.05.15.654236

**Authors:** Christian Much, Sandy M. Rajkumar, Liming Chen, John M. Cohen, Aravind R. Gade, Geoffrey S. Pitt, Yicheng Long

## Abstract

The dynamic regulation of epigenetic states relies on complex macromolecular interactions. PRC2, the methyltransferase complex responsible for depositing H3K27me3, interacts with distinct accessory proteins to form the mutually exclusive subcomplexes PHF1-PRC2.1, MTF2-PRC2.1, PHF19-PRC2.1, and PRC2.2. The functions of these subcomplexes are unclear and thought to be highly redundant. Here we show that PRC2 subcomplexes have distinct roles in epigenetic repression of lineage-specific genes and stem cell differentiation. Using a human pluripotent stem cell model, we engineered a comprehensive set of separation-of-function mutants to dissect the roles of individual protein-protein and DNA-protein interactions. Our results show that PRC2.1 and PRC2.2 deposit H3K27me3 locus-specifically, resulting in opposing outcomes in cardiomyocyte differentiation. We find that MTF2 stimulates PRC2.1-mediated repression in stem cells and cardiac differentiation through its interaction with DNA and H3K36me3, while PHF19 antagonizes it. Furthermore, MTF2-PRC2.1 maintains normal cardiomyocyte function. Together, these results reveal the importance and specificity of individual macromolecular interactions in Polycomb-mediated epigenetic repression in human stem cells and differentiation.

**Highlights:**

- The PRC2.1 and PRC2.2 subcomplexes have distinct specificities for H3K27me3 deposition
- The three PCL accessory proteins have distinct functions in regulating the PRC2 core complex, with MTF2 and PHF19 antagonizing each other
- Interactions between the PCL proteins and the PRC2 core, DNA, and H3K36me3 dictate PRC2 occupancy and activity at developmental genes
- PRC2.1 and PRC2.2 play opposing roles in stem cell cardiomyocyte differentiation
- MTF2 plays key functions in regulating differentiation timing and action potential rhythm in cardiomyocytes

## Introduction

The dynamic regulation of the epigenetic landscape is essential for the precise control of gene expression during body development.^1^ Trimethylation of lysine 27 on histone H3 (H3K27me3) is the hallmark of facultative heterochromatin and dynamically controls epigenetic repression throughout development.^2^ During embryonic development, H3K27me3 represses non-lineage genes, while removal of H3K27me3 primes lineage-specific genes for activation.^3, 4^

The sole writer complex of H3K27me3 is Polycomb Repressive Complex 2 (PRC2), which catalyzes the mono-, di-, and trimethylation of H3K27.^3^ The PRC2 core complex consists of four subunits: EZH1/2 (the catalytic subunit), SUZ12, EED, and RBBP4. PRC2 has been shown to be an essential regulator of cell differentiation and cell homeostasis, and dysregulation of PRC2 frequently leads to human diseases such as cancer and developmental defects.^5–7^ Targeting dysregulated PRC2 with small molecule inhibitors has proven effective in treating human diseases such as cancer.^8–10^ A major obstacle to fine-tuning (rather than globally inhibiting) PRC2-mediated repression in future therapy is the unclear understanding of how its activities and targeting specificity are dynamically regulated in a spatiotemporal manner.

How PRC2 is recruited to specific sites on chromatin to dynamically control epigenetic repression in mammals remains an unanswered question. Recent reports have shown that PRC2’s accessory proteins play key roles in the regulation of PRC2.^4, 11–14^ Depending on which accessory proteins the PRC2 core complex is associated with, two major PRC2 subcomplexes exist in the cell: PRC2.1, which contains one of the three Polycomb-like (PCL) proteins PHF1, MTF2, or PHF19 and occasionally EPOP and PALI1/2; and PRC2.2, which contains AEBP2 and JARID2 (reviewed in ^15^). These accessory proteins contain functional domains that interact with DNA, RNA, the PRC2 core complex, as well as other epigenetic modifications such as H2AK119ub (AEBP2 and JARID2^16^) and H3K36me3 (PCL proteins^17–19^). It remains unclear whether these dynamic macromolecular interactions have overlapping or distinct roles in epigenetic repression.

Here we show that the specificity of PRC2-mediated epigenetic repression is regulated by a set of macromolecular interactions: the interaction between the PRC2 core complex and different PRC2 accessory proteins, and interactions between individual PRC2 accessory proteins and chromatin features such as histone modifications and DNA sequences. Instead of relying solely on the conventional depletion of the entire protein or its domains, we performed “surgical operations” to engineer separation-of-function mutations to precisely disrupt specific individual macromolecular interactions. Using these separation-of-function mutants, we identified distinct roles of the two PRC2 subcomplexes PRC2.1 and PRC2.2 in H3K27me3 deposition and opposing roles in stem cell cardiac differentiation. We further determined the distinct roles of the three PRC2.1 subcomplexes containing each of the three PCL proteins using a combination of gene depletion and separation-of-function approaches. We subsequently identified distinct roles of PCL protein interactions with the PRC2 core complex, chromatin DNA, and H3K36me3 histone modification in epigenetic repression. We propose that all these dynamic macromolecular interactions together contribute to fine-tuning the epigenetic state of developmental genes during stem cell differentiation.

## Results

### Distinct roles of the PRC2.1 and PRC2.2 subcomplexes in H3K27me3 deposition

The SUZ12 subunit of the PRC2 core complex has been shown to be an important interaction hub with multiple PRC2 accessory proteins.^13, 16, 20^ Two separation-of-function mutants of SUZ12 have recently been validated in human stem cells to separate the PRC2.1 and PRC2.2 subcomplexes: loss-of-PRC2.1 mutant ((338-353)-to-GSGSGS) and loss-of-PRC2.2 mutant (F99A, R103A, L105A, I106A) of SUZ12 to yield PRC2.2-only and PRC2.1-only complexes, respectively (**Figure 1A**).^13^ The identification of these two SUZ12 mutants led to a prime opportunity to study the roles of the two subcomplexes, which in polyclonal cell lines were found to have opposite changes in SUZ12 occupancy.^13^ To further explore changes in H3K27me3 distribution, transcriptome and cell differentiation, here we derived homozygous WT control, loss-of-PRC2.1 mutant SUZ12, and loss-of-PRC2.2 mutant SUZ12 clones from the human induced pluripotent stem cell (hiPSC) line WTC-11 using a dual antibiotic selection strategy for dual-allelic genome editing (**Figure S1A**). Genomic PCR and Sanger sequencing confirmed the homozygous state as well as the correct mutations in the two mutant lines (**Figures S1B and S1C**). The edited WT and mutant SUZ12 proteins contain a 3XFLAG tag and were expressed at similar levels as validated by western blotting (**Figures 1B and S1D**) and immunofluorescence staining (**Figures 1C and S1E**). Immunofluorescence staining for OCT4 further confirmed that all three lines remained pluripotent (**Figure S1E**).

**Figure 1.**
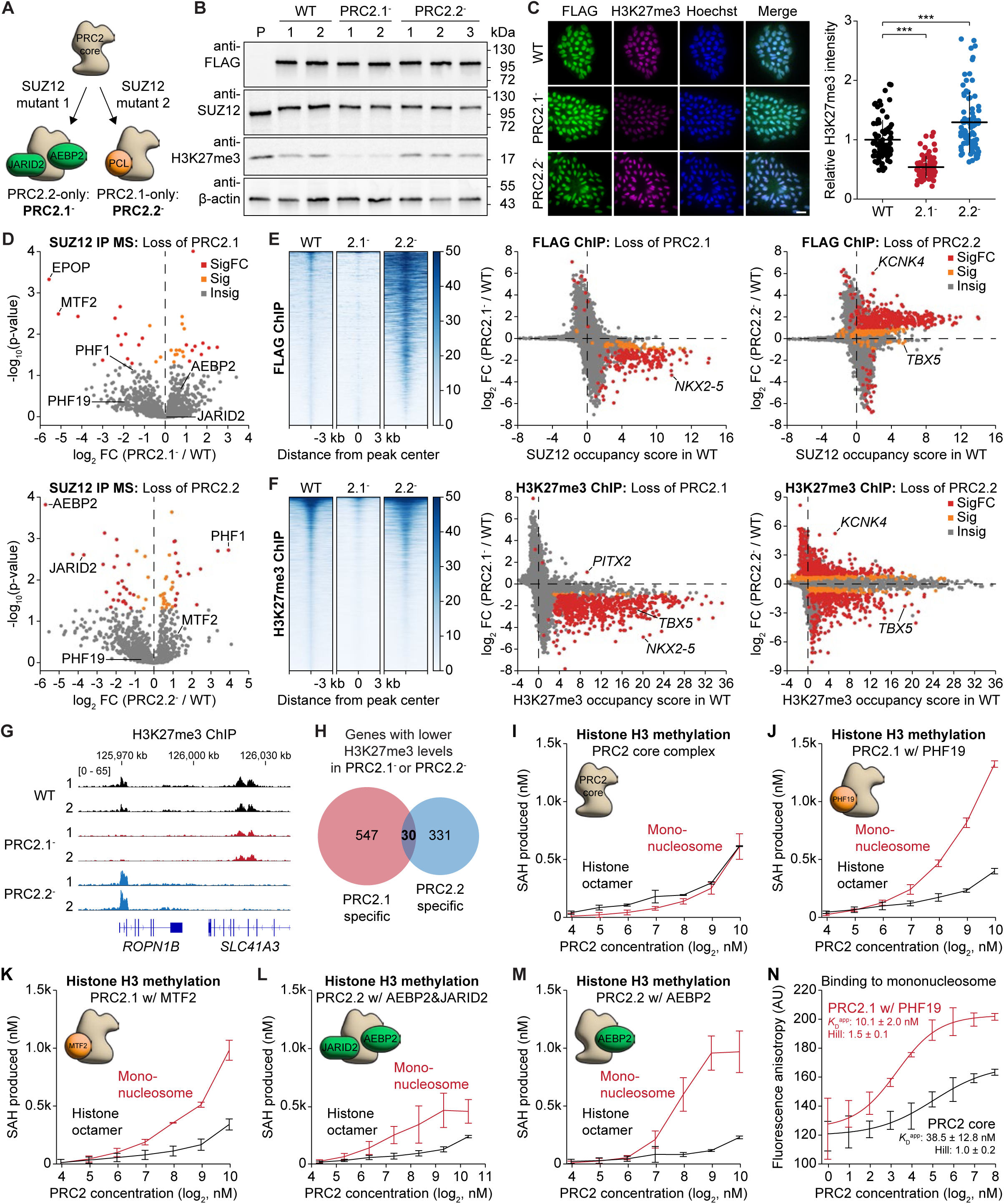
Distinct roles of the PRC2.1 and PRC2.2 subcomplexes in H3K27me3 deposition. (A) Schematic showing the separation-of-function mutants to split the PRC2.1 and PRC2.2 subcomplexes in hiPSCs. (B) Western blot on whole-cell lysates of CRISPR-edited homozygous 3XFLAG-tagged WT and the mutant SUZ12 hiPSCs PRC2.1^-^ and PRC2.2^-^. Numbers on top indicate the independent clones for each line, P denotes the untagged parental line. (C) Immunofluorescence micrographs of WT and mutant SUZ12 hiPSC lines stained with anti-FLAG (green) and anti-H3K27me3 antibody (magenta). Nuclei are stained with Hoechst (blue). Scale bar represents 20 µm. Quantification of the H3K27me3 signal is shown on the right as mean and standard deviation of 80 quantified cells each. *** *P* < 0.001, two-tailed Student’s *t*-test. (D) Mass spectrometry analysis of proteins co-immunoprecipitating with SUZ12 in WT and mutant SUZ12 hiPSC lines. Changes in the proteome in the PRC2.1^-^ (top, n = 2) and PRC2.2^-^ line (bottom, n = 3) compared to the WT (n = 2) line are shown. PRC2 accessory proteins are marked. Grey dots (Insig) indicate a *p*-value ≥ 0.05, orange dots (Sig) a *p*-value < 0.05 and |log2FC| < 1, and red dots (SigFC) a *p*-value < 0.05 and |log2FC| ≥ 1. (**E** and **F**) ChIP-seq analysis comparing genome-wide chromatin occupancy changes of FLAG-tagged SUZ12 (E) and H3K27me3 (F) upon disruption of PRC2.1 or PRC2.2. Heatmaps with SUZ12 and H3K27me3 peaks centered and sorted by decreasing peak signal based on the WT peak profile are shown on the left. Gene scatter plots depicting the enrichment score of SUZ12 or H3K27me3 in the WT and the fold change of enrichment in the mutant lines over the WT are shown on the right. Two independent clones of each genotype (n = 2) were compared by empirical Wald tests for individual genes. Grey dots (Insig) indicate a multiple test-corrected (FDR) *p*-value ≥ 0.1, orange dots (Sig) an FDR-adjusted *p*-value < 0.1 and |log2FC| < 1, and red dots (SigFC) an FDR-adjusted *p*-value < 0.1 and |log2FC| ≥ 1. (**G**) H3K27me3 ChIP-seq genome tracks showing the two adjacent genes *ROPN1B* and *SLC41A3* that have distinct PRC2 subcomplex specificity requirements. (**H**) Venn diagram showing the overlap between PRC2.1 and PRC2.2-repressed genes as defined by their loss of H3K27me3 (Sig or SigFC in (F), log2FC(mut/WT)<0, and significantly enriched between IP and input in the WT line) upon PRC2.1 or PRC2.2 disruption. (**I**-**M**) Histone H3 methylation assay using recombinant PRC2 subcomplexes on either reconstituted mononucleosome (red curves) or histone octamer alone (black curves). Data are shown as mean and standard deviation of three independent experiments. (**N**) Fluorescence polarization assay measuring the binding affinity between the PRC2 core or the PHF19-containing PRC2.1 complex and fluorescently labeled mononucleosome. Error bars represent the standard deviation of three independent experiments.

We next performed immunoprecipitation of SUZ12 followed by mass spectrometry analysis (IP-MS) to confirm the specific loss of PRC2.1 or PRC2.2-associated proteins in each respective mutant. As expected, we found that the PRC2.1-associated accessory proteins MTF2 and EPOP were the most dissociated proteins in the loss-of-PRC2.1 mutant (35- and 48-fold less enriched, respectively), while the PRC2.2-associated accessory proteins AEBP2 and JARID2 were among the most dissociated proteins in the loss-of-PRC2.2 mutant (53- and 13-fold less enriched, respectively) (**Figures 1D, S2A, and S2B**). We also identified new PRC2.1- and PRC2.2-associated proteins (**Figure S2C, and Table S1**). While KIN, HDLBP, FAU, ZSCAN10, LIG3, EIF3D, GTPBP4, EIF3A, and EIF4G1 were PRC2.1-enriched, PELP1 and RTRAF were found to be specifically enriched in PRC2.2. Interestingly, several proteins interacted with both PRC2.1 and PRC2.2 subcomplexes more strongly than with the PRC2 core complex, including RNA-binding protein G3BP2, FACT complex protein SUPT16H, transcription initiation factor TAF3, and chromatin binding protein BAP18. Additionally, we identified novel gains of interaction upon disruption of PRC2.1 or PRC2.2. For example, the H3K9 methyltransferase EHMT2 was fourfold more enriched with SUZ12 when PRC2.1 was disrupted, suggesting a role of PRC2.1 in preventing unintended epigenetic crosstalk. On the other hand, PHF1, one of the three PCL proteins in the PRC2.1 subcomplex, was 15-fold more enriched with SUZ12 upon disruption of PRC2.2 (Figures 1D and S2B), implicating that loss of PRC2.2 may result in the formation of more PHF1-containing PRC2.1 subcomplexes.

We next wanted to assess H3K27me3 levels in the SUZ12 variant lines. Western blot and immunofluorescence analyses showed that while SUZ12 levels were similar between WT and SUZ12 mutants, H3K27me3 levels were decreased in the loss-of-PRC2.1 line but increased in the loss-of-PRC2.2 line compared to WT levels, indicating different effects of PRC2.1 and PRC2.2 in H3K27me3 deposition (**Figures 1B, 1C, S1D, and S1E**). We then asked whether the perturbation of PRC2 regulation and H3K27me3 deposition was locus specific. We therefore performed FLAG and H3K27me3 ChIP-seq on the SUZ12 variant lines. FLAG ChIP-seq results showed that perturbation of PRC2.1 evicted SUZ12 from chromatin while perturbation of PRC2.2 increased SUZ12 chromatin occupancy (**Figures 1E**, **S2D, and S2H**), consistent with what was reported previously.^13^ Beyond that, H3K27me3 ChIP-seq analysis revealed that perturbation of PRC2.1 abolished H3K27me3 peaks both globally and on most PRC2 target genes (**Figures 1F and S2H, and Table S2**), consistent with the global loss of PRC2 core complex occupancy (**Figure 1E**). Interestingly, perturbation of PRC2.2 led to a bidirectional change in H3K27me3 levels (**Figure 1F and Table S3**), suggesting that PRC2.2 plays both inhibitory and stimulating roles in H3K27me3 deposition.

When examining the change in H3K27me3 levels more closely, we observed distinct H3K27 methylation specificity among genomic loci. For example, the adjacent *ROPN1B* and *SLC41A3* genes exhibit comparable H3K27me3 levels in WT hiPSCs, but specifically require PRC2.1 and PRC2.2 for H3K27me3 deposition, respectively (**Figure 1G**). Loss of PRC2.2 also resulted in an increased H3K27me3 level on *ROPN1B*, which could be directly caused by the gain of interaction between the PCL proteins and the PRC2 core complex (IP-MS data in **Figures 1D and S2B**). In contrast, loss of PRC2.1 did not increase PRC2.2-mediated H3K27me3 deposition at the *SLC41A3* gene locus, consistent with the unchanged SUZ12-JARID2/AEBP2 interaction observed in the SUZ12 IP-MS (**Figure S2B**).

This gene-specific requirement for either PRC2.1 or PRC2.2 was also observed globally: loss of PRC2.1 led to a significant loss of H3K27me3 on 577 genes, while loss of PRC2.2 caused a significant loss of H3K27me3 on 361 genes. In contrast to the previous hypothesis that PRC2.1 and PRC2.2 have overlapping roles in H3K27me3 deposition,^13, 14^ we observed that H3K27me3 was co-regulated by both PRC2.1 and PRC2.2 on only 30 genes and that PRC2.1 and PRC2.2 have a distinct specificity for H3K27me3 deposition on the majority of their regulated genes (**Figures 1H and S2E**). Gene ontology analysis revealed that PRC2.1-specific H3K27me3 deposition mainly occurred on developmental genes, especially transcription factor genes (**Figure S2F**). Genes with PRC2.1-specific H3K27me3 deposition also had a significantly higher GC content compared to genes with PRC2.2-specific H3K27me3 deposition and all genes in the genome (**Figure S2G**), consistent with PRC2.1’s role in interacting with CpG islands.^20–22^ Together, these results demonstrate that PRC2.1 and PRC2.2 have distinct roles in depositing H3K27me3 in human stem cells.

### Association with accessory proteins enhances histone methylation activity of PRC2 in a DNA-dependent manner

To biochemically evaluate how the binding of PRC2 accessory proteins affects the catalytic activity of PRC2, we tested the histone methylation activities of a series of purified recombinant PRC2 complexes on reconstituted mononucleosomes and DNA-free histone octamers (**Figure S2I**). We found that the PRC2 core complex exhibited almost identical methylation activity on mononucleosome and histone octamer substrates, suggesting that the presence of DNA in the substrate did not enhance its catalytic activity (**Figure 1I**). In contrast, association with any of the accessory proteins (PHF19, MTF2, AEBP2, and JARID2) significantly enhanced the catalytic activity of PRC2 specifically on the mononucleosome but not on the DNA-free histone octamer substrate (**Figures 1J-M**). PRC2.1 complexes exhibited higher methylation activities on the mononucleosome substrate than PRC2.2 complexes, especially at high enzyme concentrations. The stimulation of PRC2 catalytic activity on mononucleosomes is potentially caused by an increase of nucleosome-PRC2 binding affinity (**Figure 1N**). These results suggest that accessory proteins help to engage the PRC2 enzyme with the histone H3 substrate in a DNA-dependent manner.

### PRC2.1 and PRC2.2 subcomplexes distinctly regulate PRC1 recruitment and the transcriptome

PRC2 is known to crosstalk with other epigenetic modifications, such as H2AK119ub, H3K4me3, and H3K36me3.^23^ For this reason, we set out to investigate the roles of the PRC2.1 and PRC2.2 subcomplexes in regulating the epigenome beyond H3K27me3.

The PRC2 catalytic product H3K27me3 recruits PRC1 components to chromatin,^24, 25^ while the PRC2.2 components JARID2 and AEBP2 interact with the PRC1-catalyzed H2AK119ub mark.^16^ Of the two catalytic subunits of PRC1, RING1B is more highly expressed than RING1A in hiPSCs (56 TPM for RING1B vs 11 TPM for RING1A). To understand how PRC1 dynamics are affected upon perturbation of PRC2.1 and PRC2.2, we performed RING1B ChIP-seq in the loss-of-PRC2.1 and PRC2.2 lines. Perturbation of PRC2.1 led to the loss of RING1B binding to 155 genes and gain of binding to nine genes, while perturbation of PRC2.2 evicted RING1B from 176 genes and recruited it to 54 genes (**Figures 2A and S3A**). Recently, CBX7, a component of some PRC1 subcomplexes, has been shown to mediate crosstalk between JARID2-PRC2.2 and PRC1.^11^ Our CBX7 ChIP-seq indeed showed that perturbation of PRC2.2 caused a loss of CBX7 chromatin occupancy on 13 genes (**Figures 2B and S3B**). Surprisingly, perturbation of PRC2.1 evicted CBX7 from even more genes (92), which may be caused by the loss of H3K27me3 rather than the previously proposed JARID2-mediated CBX7 recruitment mechanism.^11^ In contrast to the drastic alterations of RING1B/PRC1 occupancy upon loss of PRC2.1 or PRC2.2, the genome-wide changes in H2AK119ub levels were limited to a small number of genes (four genes with lower H2AK119ub levels upon loss of PRC2.1, and five genes with lower and 13 genes with higher H2AK119ub levels upon loss of PRC2.2) (**Figure 2C**), indicating that changes in RING1B/PRC1 occupancy may not necessarily lead to subsequent changes in its catalytic product H2AK119ub.

**Figure 2.**
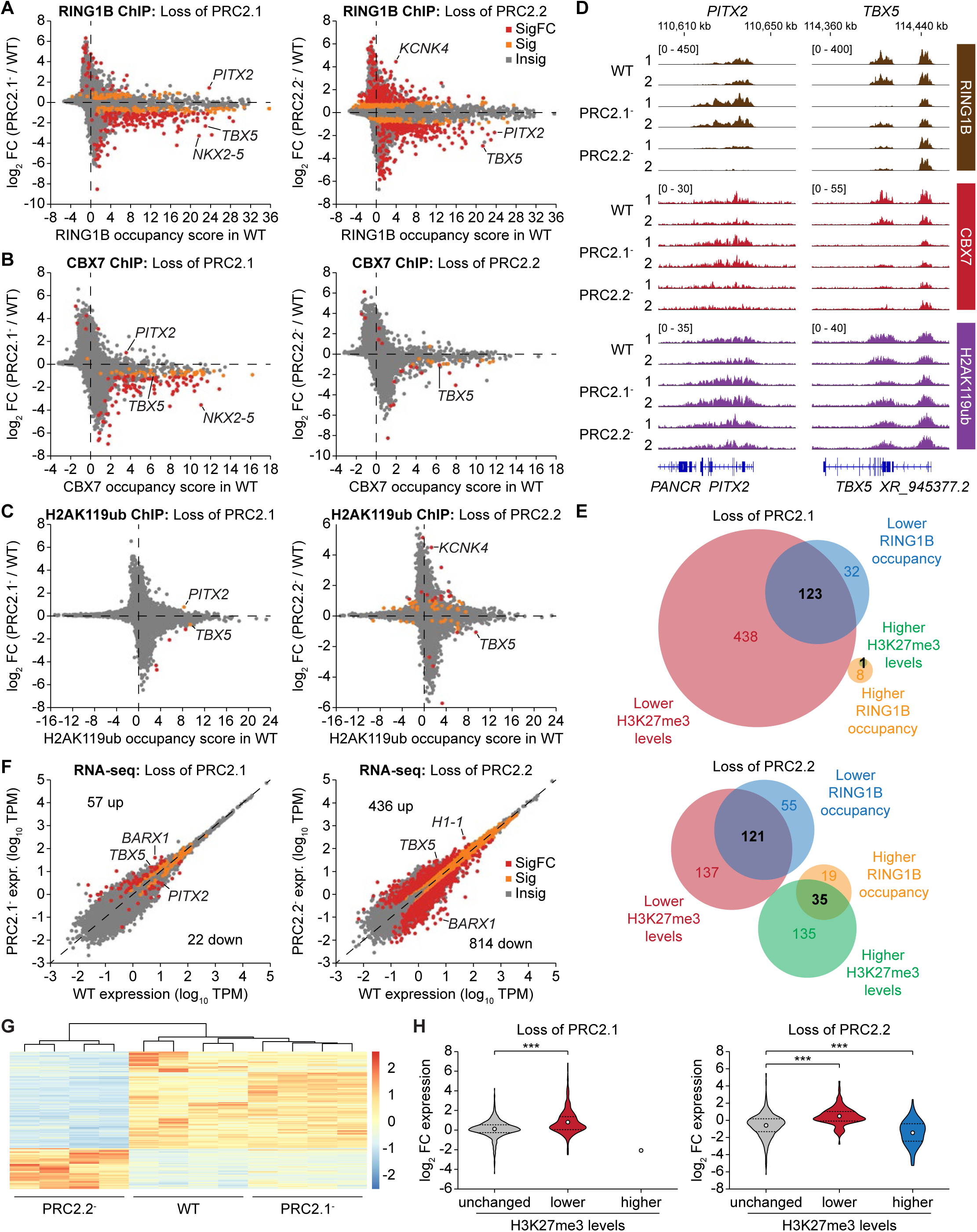
PRC2.1 and PRC2.2 subcomplexes distinctly regulate epigenome and transcriptome. (**A-C**) ChIP-seq gene scatter plots comparing genome-wide chromatin occupancy changes of RING1B (A), CBX7 (B), and H2AK119ub (C) upon disruption of PRC2.1 or PRC2.2. Two independent clones of each genotype (n = 2) were compared by empirical Wald tests for individual genes. Grey dots (Insig) indicate a multiple test-corrected (FDR) *p*-value ≥ 0.1, orange dots (Sig) an FDR-adjusted *p*-value < 0.1 and |log2FC| < 1, and red dots (SigFC) an FDR-adjusted *p*-value < 0.1 and |log2FC| ≥ 1. (D) RING1B, CBX7, and H2AK119ub ChIP-seq genome tracks of the WT and mutant SUZ12 hiPSC lines at the *PITX2* and *TBX5* gene loci. (E) Venn diagrams showing the overlap of genes with significantly altered H3K27me3 levels and RING1B occupancy upon loss of PRC2.1 or PRC2.2. (F) RNA-seq scatter plots showing gene expression changes upon loss of PRC2.1 or PRC2.2 in hiPSCs. Two independent clones of each genotype cultured independently twice (n = 4) were compared for statistical significance (two-sided Wald test). Grey dots (Insig) indicate a multiple test-corrected (FDR) *p*-value ≥ 0.05, orange dots (Sig) an FDR-adjusted *p*-value < 0.05 and |log2FC| < 1, and red dots (SigFC) an FDR-adjusted *p*-value < 0.05 and |log2FC| ≥ 1. Number of up- and downregulated genes (SigFC, red dots) are indicated. (G) Gene cluster heatmap showing the gene expression changes among WT, PRC2.1^-^ and PRC2.2^-^ hiPSC lines. All depicted genes are SigFC genes in either PRC2.1^-^ or PRC2.2^-^ lines compared to the WT (red dots in (F)). (H) Violin plots of gene expression changes upon loss of PRC2.1 (left) or PRC2.2 (right) for all H3K27me3-decorated genes in WT hiPSCs, divided into three groups based on the change of H3K27me3 levels upon depletion of the respective PRC2 subcomplex. Only one gene, *FEZF1*, had increased H3K27me3 levels upon loss of PRC2.1. *** *P* < 0.001, Welch’s *t*-test.

PRC2.1 and PRC2.2 played opposite roles in recruiting RING1B to a subset of 15 genes, including *PITX2*, a transcription factor gene that regulates muscle development (**Figure 2D**). In contrast, both PRC2.1 and PRC2.2 were required at *TBX5*, a transcription factor gene essential for the cardiac lineage (**Figure 2D**). The change in RING1B/PRC1 chromatin occupancy mimicked the change in H3K27me3 levels upon disruption of PRC2.1 or PRC2.2 (**Figure 2E**). This result supports the previous model that H3K27me3 levels determine RING1B/PRC1 occupancy on chromatin, which also agrees with recent high-throughput proteomics results showing that RING1B specifically recognizes H3K27me3-modified nucleosomes.^26^

To survey how perturbation of the PRC2 subcomplexes affects other epigenetic modifications, we performed ChIP-seq on H3K4me3 and H3K36me3, both of which have been shown to antagonize PRC2 activity.^27–29^ We found that loss of either PRC2.1 or PRC2.2 resulted in changes of H3K4me3 or H3K36me3 on a relatively small group of genes, with the loss of PRC2.2 impacting more genes compared to the loss of PRC2.1 (**Figures S3C and S3D**).

We then performed RNA-seq to compare the transcriptomic impact upon loss of PRC2.1 and PRC2.2 (**Figures 2F and 2G**). Loss of PRC2.1 resulted in an upregulation of PRC2.1-repressed genes, most of which are developmental transcription factor genes and key signaling protein genes (**Figure S3E and Table S4**). The change in RNA transcript abundance anti-correlated with the change in H3K27me3 levels for each gene decorated with H3K27me3 (**Figure 2H**). Meanwhile, we observed a much larger transcriptomic change in the loss-of-PRC2.2 lines (**Figures 2F and 2G, and Table S5**), consistent with the drastic alteration of H3K27me3 levels and PRC1/2 chromatin occupancy in this cell line. In summary, we show that loss of PRC2.1 and PRC2.2 causes distinct changes to the epigenetic landscape such as PRC1 recruitment and transcriptomic responses in undifferentiated human stem cells.

### PRC2.2 promotes, while PRC2.1 antagonizes cardiomyocyte differentiation

We next investigated the importance of PRC2.1 and PRC2.2 subcomplexes in stem cell differentiation. Notably, a previous depletion study using a degron system indicated important roles of the PRC2 subcomplexes in neural differentiation.^12^ As loss of PRC2.1 or PRC2.2 impacted the epigenome of two master regulators of cardiac differentiation (*NKX2.5* and *TBX5*, **Figures 1E and 1F**), we induced the differentiation of the WT, loss-of-PRC2.1, and loss-of-PRC2.2 lines towards cardiac progenitor cells (day 8) and cardiomyocytes (day 15) using a previously established protocol (**Figure 3A**).^30, 31^ We observed that perturbation of PRC2.1 and PRC2.2 led to distinct and opposing effects on cardiac differentiation. The loss-of-PRC2.1 line exhibited accelerated differentiation as robust spontaneous contractions were observed by day 8 (**Figures 3B and 3C**). In contrast, the loss-of-PRC2.2 line showed delayed and weaker spontaneous contractions. These opposing roles in cardiac differentiation were further reflected by the fraction of cells expressing cardiac troponin T (cTnT, a cardiac sarcomere protein and cardiomyocyte marker) as determined by flow cytometry: Perturbation of PRC2.1 increased, and perturbation of PRC2.2 decreased, the percentage of cTnT-expressing cells compared to the WT (**Figures 3D and S4A**). Immunofluorescence imaging of day 15 cardiomyocytes additionally showed that the loss-of-PRC2.2 line exhibited a lower percentage of cells expressing the cardiac marker genes alpha-actinin (*ACTN2*) and *TBX5* (**Figures 3E and S4B**). ACTN2-expressing cardiomyocytes also featured decreased H3K27me3 levels in the loss-of-PRC2.1 line (**Figure 3E**), similar to the H3K27me3 dysregulation observed in the undifferentiated hiPSC lines (**Figures 1B, 1C, and 1F**).

**Figure 3.**
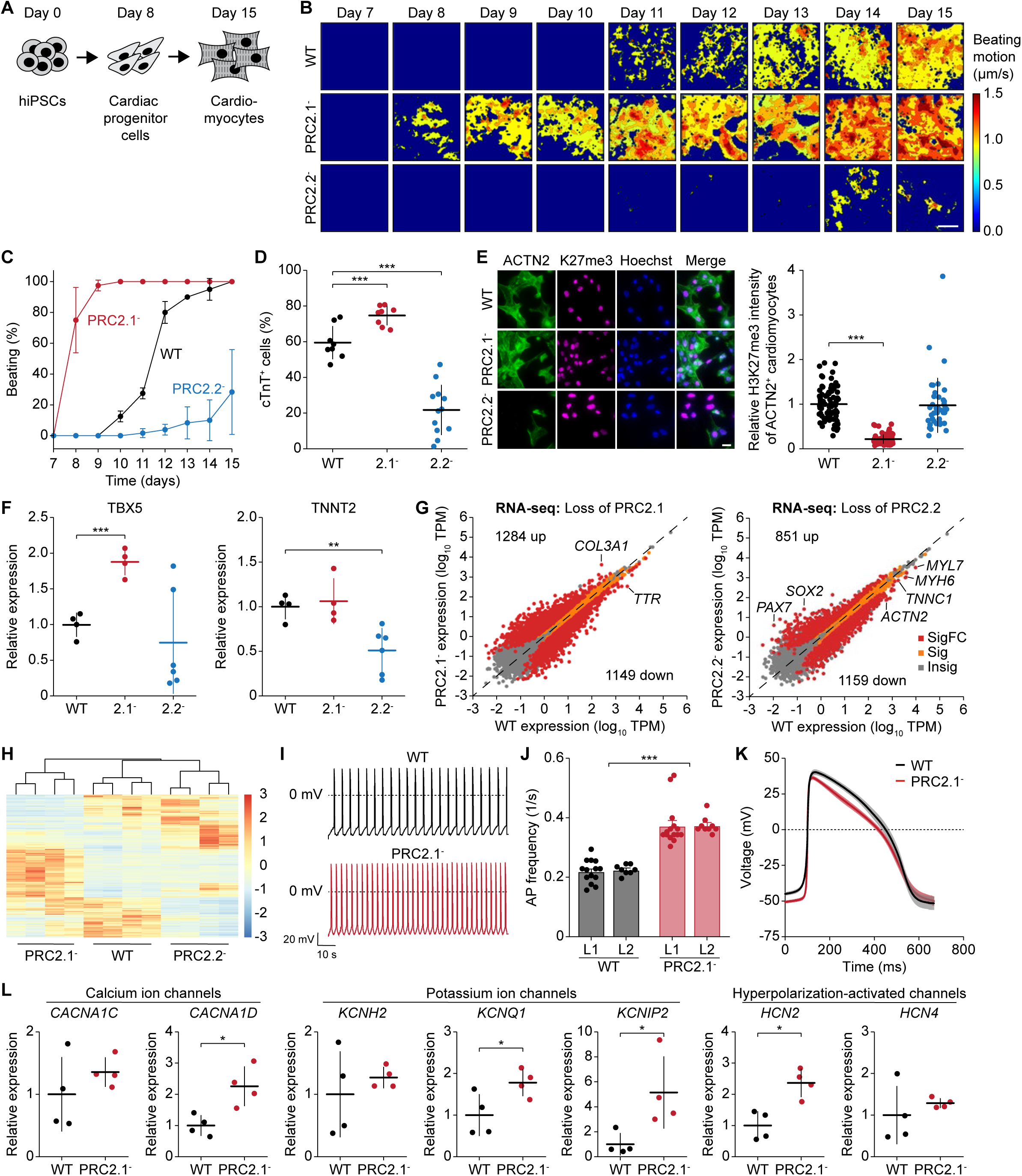
Distinct functions of the PRC2.1 and PRC2.2 subcomplexes in stem cell cardiomyocyte differentiation. (A) Schematic showing the induced cardiac differentiation of hiPSC to day 8 cardiac progenitor cells and day 15 cardiomyocytes. (B) Heatmaps showing the time-averaged magnitude of spontaneous contractions of WT, PRC2.1^-^, and PRC2.2^-^ cardiomyocytes along the differentiation process. Each heatmap is representative of eight independent recordings. (C) Percentage of cell culture wells with spontaneous contractions of WT and mutant SUZ12 cardiomyocytes over time. Data are shown as mean and standard deviation of independent clones of each genotype (n = 2 of 40 wells for WT, n = 2 of 40 wells for PRC2.1^-^, and n = 3 of 50 wells for PRC2.2^-^). (D) Percentage of WT and mutant SUZ12 cells expressing cTnT after 15 days of differentiation as determined by flow cytometry. Data are shown as mean and standard deviation of each genotype (n = 8 for WT, n = 8 for PRC2.1^-^, and n = 12 for PRC2.2^-^). *** *P* < 0.001, two-tailed Student’s *t*-test. (E) Immunofluorescence micrographs of WT and mutant SUZ12 day 15 cardiomyocytes stained with anti-ACTN2 (green) and anti-H3K27me3 antibody (magenta). Nuclei are stained with Hoechst (blue). Scale bar represents 20 µm. Quantification of the H3K27me3 signal of ACTN2^+^ day 15 cardiomyocytes are shown on the right as mean and standard deviation of 83 (WT), 97 (PRC2.1^-^), and 41 (PRC2.1^-^) quantified cells. *** *P* < 0.001, two-tailed Student’s *t*-test. (F) Relative expression of cardiomyocyte marker genes in day 15 cardiomyocytes as determined by qRT-PCR analysis. Data are shown as mean and standard deviation of four (WT and PRC2.1^-^) or six (PRC2.2^-^) biological replicates each. * *P* < 0.05, ** *P* < 0.01, *** *P* < 0.001, two-tailed Student’s *t*-test. (G) RNA-seq scatter plots showing gene expression changes upon loss of PRC2.1 or PRC2.2 in day 15 cardiomyocytes. Two independent clones of each genotype cultured independently twice (n = 4) were compared for statistical significance (two-sided Wald test). Grey dots (Insig) indicate a multiple test-corrected (FDR) *p*-value ≥ 0.05, orange dots (Sig) an FDR-adjusted *p*-value < 0.05 and |log2FC| < 1, and red dots (SigFC) an FDR-adjusted *p*-value < 0.05 and |log2FC| ≥ 1. Number of up- and downregulated genes (SigFC, red dots) are indicated. (H) Gene cluster heatmap showing the gene expression changes among WT, PRC2.1^-^ and PRC2.2^-^ day 15 cardiomyocytes. All depicted genes are SigFC genes in either PRC2.1^-^ or PRC2.2^-^ lines compared to the WT (red dots in (G)). (**I-K**) Electrophysiological analysis of WT and PRC2.1-day 24 cardiomyocytes as recorded by patch clamp. Representative action potential (AP) traces (I), bar plots of average AP frequency (J), and average single AP trace of all measured cells over time (K) of spontaneous contractions are shown. Data in (J and K) are shown as mean and standard error of the mean of 14 (WT replicate 1), 8 (WT replicate 2), 15 (PRC2.1^-^ replicate 1), and 9 (PRC2.1^-^ replicate 2) cell measurements. *** *P* < 0.001, two-way ANOVA. (**L**) Relative expression of ion channel genes in day 24 cardiomyocytes as determined by qRT-PCR analysis. Data are shown as mean and standard deviation of four biological replicates each. * *P* < 0.05, two-tailed Student’s *t*-test.

We then asked about the consequences on gene expression in the SUZ12 mutant hiPSC-derived cardiomyocytes. qRT-PCR results indicate that the loss-of-PRC2.1 line featured significantly higher *TBX5* expression levels, while the loss-of-PRC2.2 line expressed significantly less *TNNT2* (**Figure 3F**). Beyond that, RNA-seq analysis revealed a massive dysregulation of gene expression in both lines (**Figures 3G and 3H, and Tables S6 and S7**), with most dysregulated genes having key roles in development (**Figure S4C**).

Next, we aimed to determine if the loss of PRC2.1 and PRC2.2 affected the development of the cardiomyocytes’ electrical properties. To this end, we enriched the cardiomyocyte population using lactate-based purification and recorded spontaneous action potentials of day 24 cardiomyocytes. We were unable to assess these properties in the loss-of-PRC2.2 line, as their global defect in cardiac differentiation yielded very few spontaneously contracting cells. In contrast, we found that the loss-of-PRC2.1 cardiomyocytes featured an action potential frequency that was almost doubled compared to the WT, which was consistent across the two independent CRISPR clones for each line (**Figures 3I and 3J**). Interestingly, there was a trend for faster depolarization in the PRC2.1^-^ line (**Figure 3K**), which is also reflected by the significantly lower action potential duration at 30% repolarization level (APD30) (**Figure S4E, left**). Other parameters of the action potential remained unchanged upon loss of PRC2.1, such as action potential amplitude and duration at 50% (APD50) or 90% (APD90) repolarization level (**Figures S4D and S4E**). The significantly increased action potential frequency points to an ion transport abnormality, potentially caused by a de-repression and upregulation of ion channel genes. While increased expression of multiple ion channels likely contributes to the increased action potential frequency, we observed increased expression of *HCN2* (potassium/sodium hyperpolarization-activated cyclic nucleotide-gated channel 2), *KCNQ1* (potassium channel Kv7.1), and *KCNIP2* (potassium voltage-gated channel interacting protein 2) upon perturbation of PRC2.1 in day 23 cardiomyocytes (**Figure 3K**).

Upregulation of *HCN2* channels would increase spontaneous pacing rates and upregulation of *KCNQ1* channels or the *KCNIP2*-encoded KChIP2 auxiliary subunit could accelerate repolarization, all of which would support an increased action potential frequency. Together, we conclude that the PRC2 subcomplexes exert distinct functions during cardiac differentiation, with PRC2.2 promoting and PRC2.1 antagonizing the process.

### Distinct and overlapping roles of PCL proteins in PRC2.1 function

PRC2.1 can be further divided into three mutually exclusive subcomplexes that are formed through the interaction between the core complex and one of the three PCL proteins – PHF1, MTF2, or PHF19 (**Figure 4A**). Among the three PCL proteins, MTF2 shows the highest expression at the RNA level in hiPSCs (**Figure 4B**), although PCL protein expression patterns can change upon differentiation.^32^ Previous studies have shown that the PCL proteins are essential for PRC2 function (reviewed in ^15^). However, it remains unclear whether the three PCL proteins have overlapping or distinct functions.

**Figure 4.**
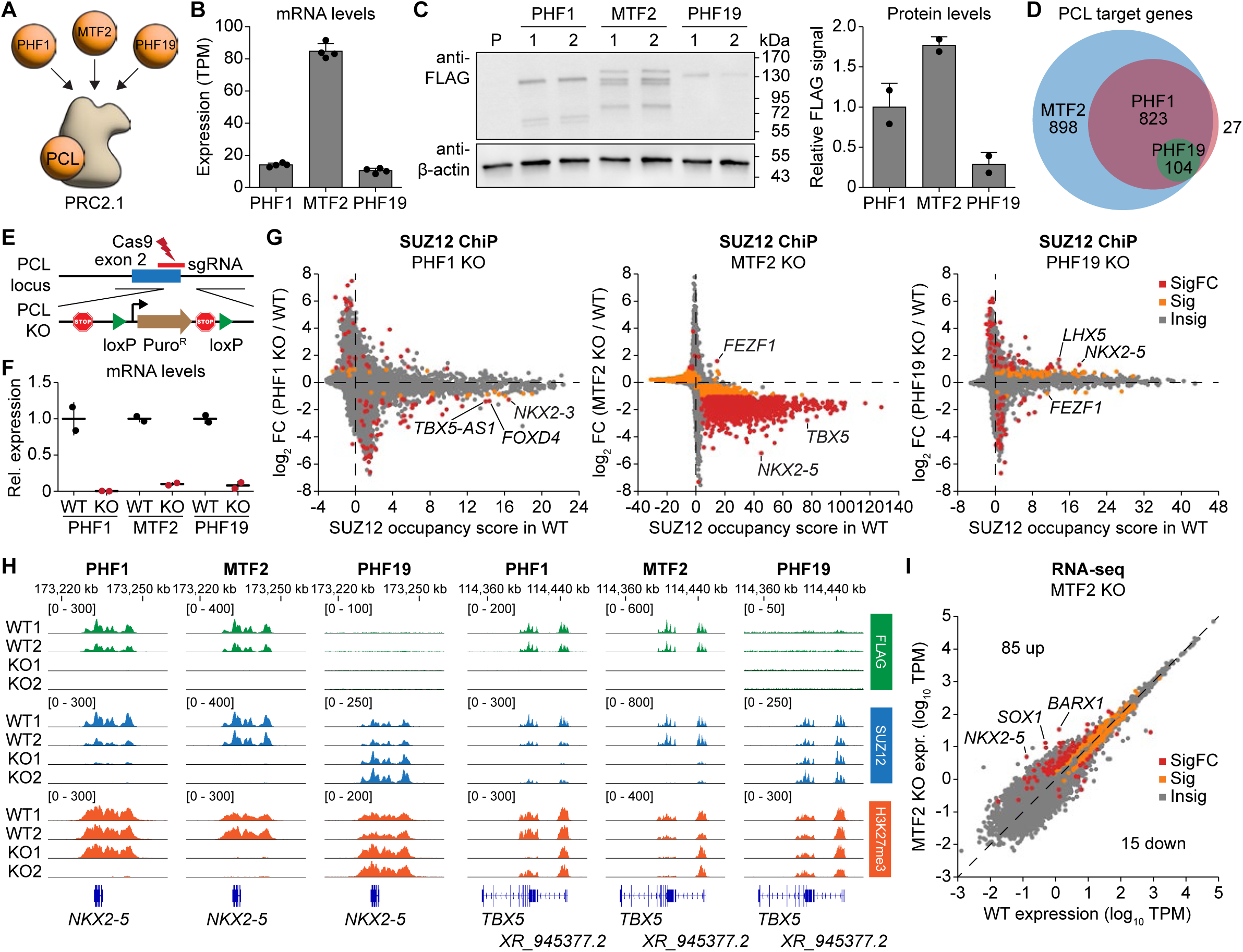
Distinct and overlapping roles of PCL proteins in PRC2.1 function. (A) Schematic illustrating the three PCL proteins PHF1, MTF2, and PHF19 associating with the PRC2 core complex to form three mutually exclusive PRC2.1 subcomplexes. (B) Expression of the three PCL genes in hiPSCs as measured by RNA-seq. Data are shown as mean and standard deviation of four biological replicates each. (C) Western blot on whole-cell lysates of CRISPR-edited homozygous 3XFLAG-HALO-tagged PHF1, MTF2, and PHF19 hiPSCs. Numbers on top indicate the independent clones for each line, P denotes the untagged parental line. Note that PHF1 and MTF2 have multiple isoforms. The bar plot on the right shows the quantification of the FLAG-tagged protein signal of each PCL protein normalized to the β-actin signal. Data are shown as mean and standard deviation of two biological replicates each. (D) Venn diagram showing the number of distinct and overlapping target genes of the three PCL proteins as defined by FLAG-tagged PCL protein occupancy in the ChIP-seq experiments. (E) Schematic illustrating the PCL gene depletion strategy by inserting poly(A) sites (indicated by stop sign) together with a *loxP*-flanked puromycin resistance gene expression cassette immediately downstream of the translation start site. (F) Relative expression of each PCL gene in WT (3XFLAG-HALO tagged) and KO hiPSC lines as determined by qRT-PCR analysis. Data are shown as mean and standard deviation of two homozygous clones each. (G) ChIP-seq gene scatter plot comparing genome-wide SUZ12 chromatin occupancy changes upon loss of PHF1, MTF2, or PHF19. Two independent clones of each genotype (n = 2) were compared by empirical Wald tests for individual genes. Grey dots (Insig) indicate a multiple test-corrected (FDR) *p*-value ≥ 0.1, orange dots (Sig) an FDR-adjusted *p*-value < 0.1 and |log2FC| < 1, and red dots (SigFC) an FDR-adjusted *p*-value < 0.1 and |log2FC| ≥ 1. (H) FLAG, SUZ12, and H3K27me3 ChIP-seq genome tracks of the PCL protein WT and KO hiPSC lines at the *NKX2-5* and *TBX5* gene loci. (I) RNA-seq scatter plots showing gene expression changes upon MTF2 depletion. Two independent clones of each genotype cultured independently twice (n = 4) were compared for statistical significance (two-sided Wald test). Grey dots (Insig) indicate a multiple test-corrected (FDR) *p*-value ≥ 0.05, orange dots (Sig) an FDR-adjusted *p*-value < 0.05 and |log2FC| < 1, and red dots (SigFC) an FDR-adjusted *p*-value < 0.05 and |log2FC| ≥ 1. Number of up- and downregulated genes (SigFC, red dots) are indicated.

The difficulty in studying the individual specificity of the PCL proteins arises partly due to a lack of specific antibodies for PHF1 and PHF19. To overcome this limitation and to enable a direct comparison between the PCL proteins, we generated three hiPSC lines by attaching a 3XFLAG-HALO tag to the C-terminus of each PCL protein. Western blotting using an anti-FLAG antibody in the homozygous FLAG-tagged PCL protein lines suggests that MTF2 is the most abundant PCL protein in undifferentiated hiPSCs, followed by PHF1 and PHF19 (**Figure 4C**). Notably, multiple protein bands were observed for PHF1 and MTF2, suggesting different isoforms. To determine the chromatin targeting specificity of the PCL proteins, we performed FLAG ChIP-seq in the three edited hiPSC lines. We observed that the number of target genes – as defined by significant FLAG-tagged PCL protein chromatin occupancy – positively correlates with the protein expression level of each PCL protein (**Figure 4D**). Interestingly, their target genes are highly overlapping: all 104 PHF19 target genes are also targeted by PHF1 and MTF2, and almost all PHF1 target genes are bound by MTF2 as well (927 out of 954 genes, **Figure 4D and Table S8**). Gene ontology analysis revealed similar pathway genes in cell differentiation and transcriptional regulation for all PCL protein target genes (**Figure S5A**).

We next asked whether such high degree of shared target genes also translates to high functional redundancy. We therefore generated knockout hiPSC lines for each PCL protein by inserting polyadenylation sites downstream of their translation start sites (**Figure 4E**). Depletion of each endogenous PCL gene at the mRNA and protein level was confirmed by qRT-PCR and western blotting, respectively **(Figures 4F and S5B)**. To test how depletion of each PCL protein impacted PRC2 recruitment, we performed SUZ12 ChIP-seq comparing each PCL protein WT and KO hiPSC line. Depletion of MTF2 led to a global loss of SUZ12 chromatin occupancy, while depletion of PHF1 or PHF19 resulted in milder changes in SUZ12 occupancy (**Figures 4G and S5C**), likely reflecting PCL protein expression levels and therefore specific PCL protein-containing PRC2.1 subcomplex ratios. Two of the genes with the highest loss of SUZ12 occupancy upon depletion of MTF2 and PHF1 were the cardiac transcription factor genes *NKX2-5* and *TBX5* (**Figure 4H**). Surprisingly, depletion of PHF19 led to a gain of SUZ12 occupancy on *NKX2-5* but not *TBX5* (**Figure 4H**), suggesting that PHF19 inhibits PRC2 recruitment at selected gene loci. The altered SUZ12 occupancy resulted in more modest and gene-specific changes in H3K27me3 levels in the MTF2, PHF1, and PHF19 KO, indicating that PRC2 catalytic activity may not always positively correlate with its chromatin occupancy (**Figures S5D and S5E**). Importantly, RNA-seq showed that genes with altered expression in the MTF2 KO hiPSCs were associated with cell differentiation and transcriptional regulation (**Figures 4I and S5F**). In summary, these results suggest that the three PCL proteins have overlapping yet distinct functions in PRC2 recruitment, with MTF2 playing the most dominant role in undifferentiated hiPSCs.

### Disrupting the interaction of individual PCL proteins with the PRC2.1 subcomplex results in different epigenetic consequences

As the complete loss of PCL proteins could lead to secondary effects independent of PRC2 function, we next aimed to engineer separation-of-function mutants to disrupt the interaction between each of the three PCL proteins and PRC2 (**Figure 5A**). A recently solved crystal structure captured two potential interfaces between SUZ12 and PHF19.^20^ Interface 1 is assembled by the interaction between two α-helices: L535, I539, and F543 of PHF19 form hydrophobic interactions with I96 and F99 of SUZ12, and an additional hydrogen bonding between Y542 of PHF19 and Q95 of SUZ12 further locks in the two helices (**Figure 5B**). Interface 2 is formed by a loop region of PHF19 and a β-sheet region of SUZ12, and three residues (R561, L571, and W574) of PHF19 are potentially key to this interaction (**Figure 5B**). R561, L571, and W574 are highly conserved in all three human PCL proteins as well as the Drosophila Pcl protein, while L535, I539, Y542, and F543 are only conserved in MTF2 and PHF19 but not PHF1.

**Figure 5.**
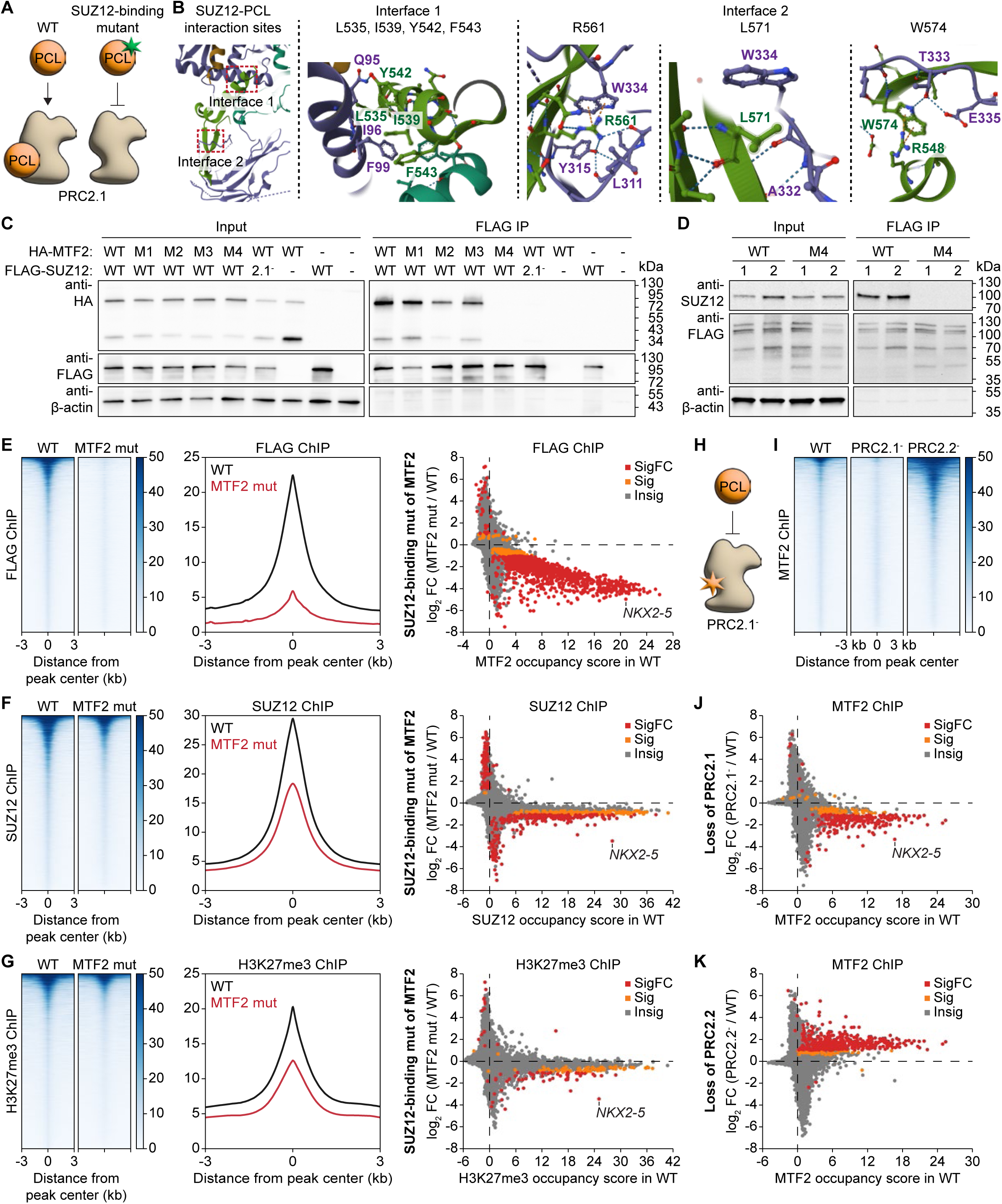
Splitting each PCL protein from the PRC2.1 subcomplex results in different epigenetic consequences. (A) Schematic showing the engineering of PCL protein separation-of-function mutants to disrupt their interaction with SUZ12 and the PRC2 core complex. (B) Crystal structure (PDB: 6NQ3) highlighting the two interaction sites between SUZ12 and PHF19. SUZ12 is depicted in purple, PHF19 in green. Amino acid residues of SUZ12 and PHF19 that are involved in the interactions are highlighted in the close-up images. The corresponding amino acid residues for MTF2 are L548, I552, Y555, F556 for interface 1 and R574, L584, W587 for interface 2. (C) Western blot of co-immunoprecipitation assay performed in HEK293T cells by transiently expressing ectopic HA-tagged MTF2 and FLAG-tagged SUZ12. Input samples represent the whole cell lysate and the FLAG-IP samples the fraction eluted off the anti-FLAG beads. Mutants of MTF2: M1: L548A, I552A, Y555A, F556A; M2: L584A, W587A; M3: R574A; M4: R574A, L584A, W587A. PRC2.1^-^ mutant of SUZ12: (338-353)GSGSGS. Note that the lower band on the anti-HA blot likely represents a cleaved product or shorter isoform of the full-length HA-MTF2 (upper band). (D) Western blot of co-immunoprecipitation assay performed in CRISRP-edited homozygous WT and SUZ12-binding mutant MTF2 hiPSCs. Whole cell lysate (input) was incubated with anti-FLAG beads to pull down the FLAG-tagged MTF2. M4 mutant as in (C): R574A, L584A, W587A. The multiple bands in the anti-FLAG Western blot represent the different MTF2 isoforms. (**E-G**) ChIP-seq analysis comparing genome-wide chromatin occupancy changes of FLAG-tagged MTF2 (E), SUZ12 (F), and H3K27me3 (G) between WT and the SUZ12-binding mutant of MTF2. Heatmaps with MTF2, SUZ12, and H3K27me3 peaks centered and sorted by decreasing peak signal based on the WT peak profile are shown on the left. Metaplots of the peak signals are shown in the middle. Gene scatter plots depicting the enrichment score of MTF2, SUZ12, and H3K27me3 in the WT and the fold change of enrichment in the mutant over the WT are shown on the right. Grey dots (Insig) indicate a multiple test-corrected (FDR) *p*-value ≥ 0.1, orange dots (Sig) an FDR-adjusted *p*-value < 0.1 and |log2FC| < 1, and red dots (SigFC) an FDR-adjusted *p*-value < 0.1 and |log2FC| ≥ 1. (H) Schematic showing the disruption of the PCL protein-PRC2 core interaction using the PRC2.1^-^ mutant. (I) Heatmaps of the MTF2 ChIP-seq peak analysis in the WT, PRC2.1^-^ and PRC2.2^-^ hiPSCs lines. (**I-K**) ChIP-seq analysis comparing genome-wide chromatin occupancy changes of MTF2 upon loss of PRC2.1 and PRC2.2. Heatmaps with MTF2 peaks centered and sorted by decreasing peak signal based on the WT peak profile are shown in (I). Gene scatter plots depicting the enrichment score of MTF2 in the WT and the fold change of enrichment in PRC2.1^-^ and PRC2.2^-^ over the WT are shown in (J) and (K), respectively. Grey dots (Insig) indicate a multiple test-corrected (FDR) *p*-value ≥ 0.1, orange dots (Sig) an FDR-adjusted *p*-value < 0.1 and |log2FC| < 1, and red dots (SigFC) an FDR-adjusted *p*-value < 0.1 and |log2FC| ≥ 1.

To pinpoint the residues that are critical for the interaction with SUZ12, we first generated a series of MTF2 mutants and used co-immunoprecipitation experiments in HEK293T cells to survey which mutations abolish the interaction between SUZ12 and MTF2. Mutation of all four residues of interface 1 on MTF2 (M1 mutant: L548A, I552A, Y555A, and F556A) did not affect the interaction with SUZ12 (**Figure 5C**). However, mutation of the three residues of interface 2 (M4 mutant: R574A, L584A, and W587A) completely abolished SUZ12 interaction. Moreover, neither the single mutant (M3 mutant: R574A) nor the double mutant (M2 mutant: L584A, W587A) was sufficient to completely disrupt the interaction (**Figure 5C**). These three conserved PCL residues of interface 2 are particularly important not only for MTF2, but also for PHF1 and PHF19, to interact with SUZ12 (**Figures S6A and S6B**).

We then introduced the identified SUZ12-binding mutations at the respective endogenous PCL gene locus in hiPSCs. Immunoprecipitation of FLAG-MTF2 followed by SUZ12 western blotting confirmed that the SUZ12 interaction was abolished in the SUZ12-binding mutant MTF2 hiPSC lines (**Figure 5D**). ChIP-seq showed that loss of the MTF2-SUZ12 interaction decreased the occupancy of both MTF2 and SUZ12 on the majority of PRC2 target genes and was concomitant with decreased H3K27me3 levels (**Figures 5E-5G**), consistent with our and previous observations in MTF2 depletion studies.^4, 11, 12, 14, 33^ Disruption of the PHF1-SUZ12 interaction led to a reduction of SUZ12 chromatin occupancy on fewer genes (74 PRC2 target genes with significantly reduced and one gene with significantly increased SUZ12 occupancy) (**Figures S6C and S6E**). In contrast, perturbation of the PHF19-SUZ12 interaction resulted in a global increase in SUZ12 chromatin occupancy (**Figures S6D and S6F**). For example, SUZ12 occupancy was increased on the *NKX2-5* gene upon loss of the PHF19-SUZ12 interaction but was decreased upon loss of the MTF2/PHF1-SUZ12 interaction. These results indicate that PHF19 antagonizes PRC2 recruitment, while MTF2 and PHF1 promote it. Our H3K27me3 ChIP-seq results further support the distinct roles of the three PCL proteins in interacting with SUZ12 (**Figures 5G, S6E, and S6F**).

Interestingly, we found that the PCL proteins depend on the PRC2 core complex to stably associate with chromatin, as FLAG ChIP-seq results demonstrated that the PCL proteins collectively exerted lower genome-wide chromatin occupancy in the SUZ12-binding mutant PCL protein hiPSC lines (**Figures 5E and S6C-S6F**). To confirm this observation further with an orthogonal approach, we performed MTF2 ChIP-seq in the loss-of-PRC2.1 and loss-of-PRC2.2 hiPSC lines (**Figure 5H)**. Perturbation of PRC2.1 led to a loss of MTF2 chromatin occupancy genome-wide (**Figures 5I and 5J**), which is in agreement with our conclusion that the PCL proteins require PRC2 to localize to chromatin. Conversely, perturbation of PRC2.2 led to increased MTF2 chromatin occupancy (**Figures 5I and 5K**), which could be caused by a gain of MTF2-PRC2 association upon loss of PRC2.2. In conclusion, we find that disrupting the interaction between the PRC2 core and each of its PCL accessory proteins results in different epigenetic outcomes, where MTF2 and PHF1 promote and PHF19 antagonizes PRC2 recruitment and H3K27me3 deposition.

### The DNA-PCL protein interaction is the main driver of Polycomb-mediated repression

In the fruit fly *Drosophila melanogaster*, PRC2 is recruited by sequence-specific DNA binding proteins that recognize Polycomb response elements.^34, 35^ In mammals, it has been shown that PRC2 can preferentially bind CpG-rich DNA sequences,^36–39^ and these DNA interactions may depend on the winged-helix domain of the PCL accessory proteins of PRC2.^20–22^ Using purified recombinant PRC2 complexes, we compared the DNA and nucleosome binding affinity of the PRC2 core complex and the MTF2-bound PRC2.1 complex. Electrophoretic mobility shift assay (EMSA) showed that the MTF2-PRC2.1 complex exhibited a higher binding affinity (*KD*app of 48 nM) to dsDNA from the *NKX2-5* promoter region than the PRC2 core complex alone (*KD*app of 116 nM) (**Figure 6A**). Significantly increased binding affinity was also observed on *HOXA9* promoter DNA, but the difference was much smaller on a non-PRC2 target *SSTR4* promoter region (**Figure S7A**). Fluorescence polarization further confirmed that binding to a reconstituted mononucleosome was significantly enhanced with the addition of MTF2 to the PRC2 core complex (**Figure 6B**).

**Figure 6.**
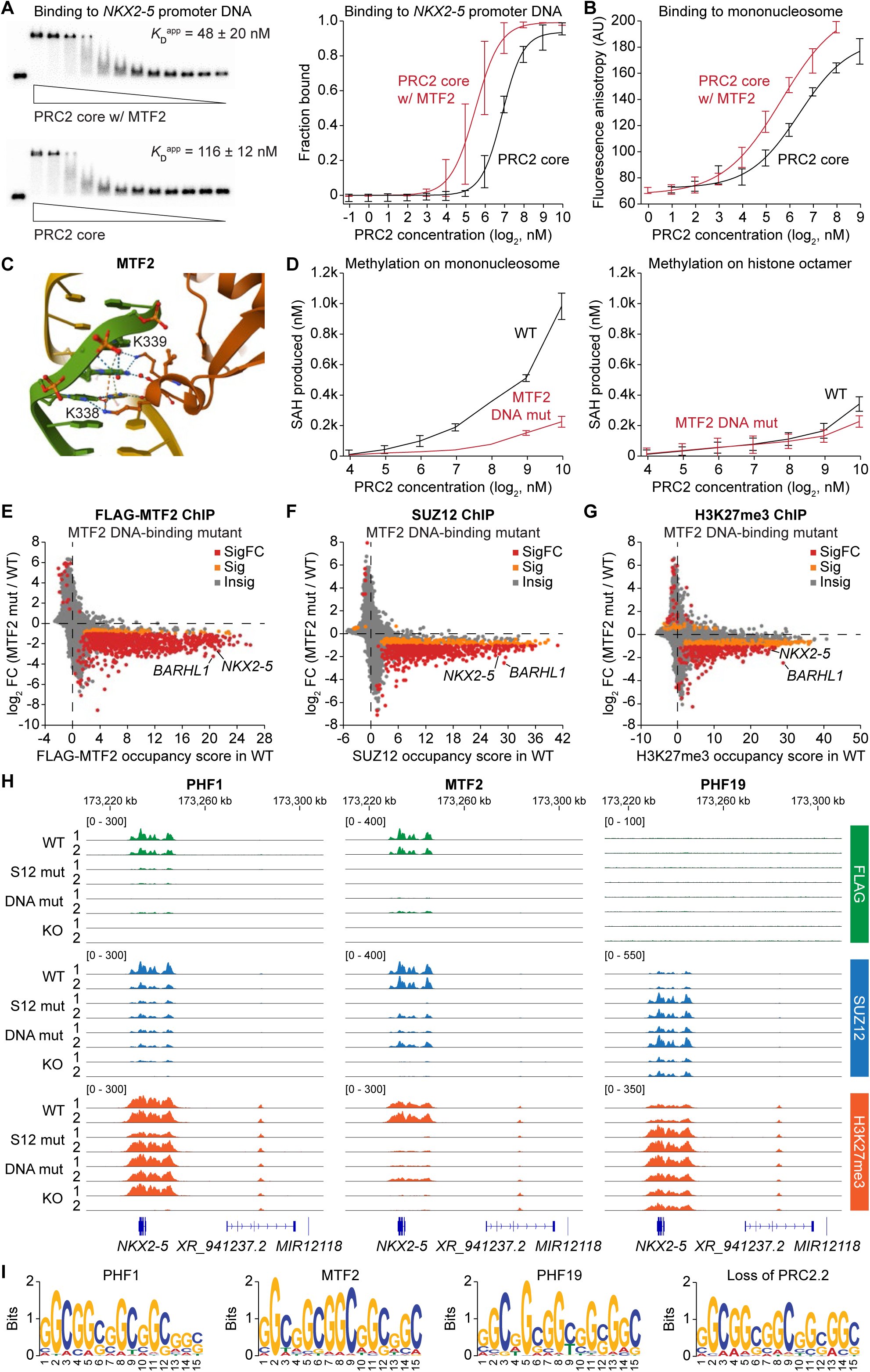
The DNA-PCL protein interaction is a main driver of PRC2-chromatin interaction. (A) EMSA to compare the binding affinity between an *NKX2-5* promoter DNA and the recombinant PRC2 core complex or the MTF2-containing PRC2.1 complex. Representative EMSA gels are shown on the left. Quantification of the fraction of DNA bound with error bars representing the standard deviation from three independent experiments is shown on the right. (B) Fluorescence polarization assay measuring the binding affinity between the PRC2 core or the MTF2-containing PRC2.1 complex and fluorescently labeled mononucleosome. Error bars represent the standard deviation of three independent experiments. (C) Crystal structure (PDB: 5XFR) highlighting the two lysine residues of MTF2 (K338 and K339) interacting with the major groove of double-stranded DNA. (D) Histone H3 methylation assay using recombinant WT or DNA-binding mutant (K338A, K339A) MTF2-containing PRC2.1 complexes on either reconstituted mononucleosome (left) or histone octamer (right). Data are shown as mean and standard deviation of three independent experiments. (**E-G**) ChIP-seq gene scatter plots comparing genome-wide chromatin occupancy changes of FLAG-MTF2 (E), SUZ12 (F), and H3K27me3 (G) between WT and DNA-binding mutant MTF2 hiPSC lines. Two independent clones of each genotype (n = 2) were compared by empirical Wald tests for individual genes. Grey dots (Insig) indicate a multiple test-corrected (FDR) *p*-value ≥ 0.1, orange dots (Sig) an FDR-adjusted *p*-value < 0.1 and |log2FC| < 1, and red dots (SigFC) an FDR-adjusted *p*-value < 0.1 and |log2FC| ≥ 1. (H) FLAG, SUZ12, and H3K27me3 ChIP-seq genome tracks of WT, SUZ12-binding mutant, DNA-binding mutant, and KO of each PCL protein around the *NKX2-5* gene locus. (I) Sequence logos of the most significant DNA motif for PHF1, MTF2, and PHF19 identified in the FLAG ChIP-seq of the respective FLAG-tagged PCL protein hiPSC lines. Note that this motif was not found in the FLAG ChIP-seq of the KO lines.

Essential amino acid residues (e.g., K338 and K339 in MTF2, K323 and K324 in PHF1, and K331 and K332 in PHF19) in the PCL protein winged-helix domain have been reported in DNA-bound crystal structures,^21^ revealing that they form hydrogen bonds with the nucleotide bases in the major groove of the dsDNA (**Figure 6C**). In order to test the importance of these residues for DNA binding and PRC2 recruitment, we generated an MTF2 mutant (K338A, K339A) in the recombinant MTF2-PRC2.1 complex. Compared to the WT PRC2.1 complex, the mutant PRC2.1 complex exhibited a significant methylation activity defect on a DNA-containing mononucleosome, while no changes in activity were observed on a histone octamer substrate alone without DNA (**Figure 6D**). This result confirmed that the two conserved lysine residues in MTF2 are critical for PRC2 engagement with the nucleosome substrate.

To investigate the importance of these residues for PRC2 function in the cell, we generated homozygous DNA-binding mutant PCL protein hiPSC lines with K338A and K339A mutations. FLAG ChIP-seq analysis revealed that both the DNA-binding mutants of PHF1 and MTF2 lost their association with chromatin (**Figures 6E and S6B**). Consequently, the DNA-binding mutant MTF2 lines exhibited a global loss of SUZ12 chromatin occupancy and decreased H3K27me3 levels on most PRC2 target genes (**Figures 6F and 6G**), while only a few genes were affected in the DNA-binding mutant of PHF1 (**Figure S7B**). Interestingly and again in contrast to MTF2, the PHF19 DNA-binding mutant showed a gain of SUZ12 occupancy and H3K27me3 levels on many PRC2-target genes (**Figure S7C**). Together with results from the SUZ12-binding PCL protein mutants, we conclude that the three PCL proteins exert distinct functions in regulating the PRC2 core complex, with MTF2 and PHF19 antagonizing each other. For example, MTF2 increased both PRC2 recruitment and H3K27me3 levels at the cardiac transcription factor gene *NKX2-5*, while PHF19 decreased them (**Figure 6H**). Of note, the distinct functions of the PCL proteins are likely not based on different DNA binding preferences, as DNA-binding motif analysis revealed that they all bind the same CGG-rich binding motif (**Figure 6I**). Moreover, this motif was also enriched in the loss-of-PRC2.2 hiPSCs, indicating that this is a common feature of PRC2.1 DNA binding (**Figure 6I**). Together, our data suggest that all three PCL proteins specifically recognize CGG-rich DNA sequences but exert distinct roles in PRC2 recruitment and H3K27me3 deposition.

### The H3K36me3-MTF2 interaction provides an alternative recruitment mechanism for PRC2 and stimulates H3K36me3 demethylation at *TBX5*

The N-terminal Tudor domain of PCL proteins interacts with H3K36me3, a histone methylation mark that frequently decorates the gene body of actively transcribed genes.^40^ It has been shown that the interaction between H3K36me3 and the Tudor domain of PHF1 and PHF19 is important for PRC2 recruitment to H3K36me3-marked genes.^17, 18^ In contrast, the Tudor domain of MTF2 has a lower affinity for H3K36me3 in biochemical binding assays,^21^ and its function remains unclear. Based on the solved structures of PHF1,^41^ MTF2,^21^ and PHF19,^18^ three residues within the Tudor domain form an aromatic cage to lock the H3K36me3 in place (**Figure 7A**). We generated an MTF2 H3K36me3-binding mutant hiPSC line by introducing the Y62A mutation. Mutation of this residue in PCL proteins (Y47A of PHF1 or Y56A of PHF19) has been validated to disrupt H3K36me3 recognition.^17, 18^ ChIP-seq on FLAG-tagged MTF2, SUZ12, and H3K27me3 revealed that the Y62A mutation on MTF2 led to a predominant decrease in MTF2 and SUZ12 chromatin occupancy and H3K27me3 levels on PRC2 target genes, including the developmental transcription factor genes *TBX5* and *BARHL1* (**Figure 7B**). These results indicate that the H3K36me3-MTF2 interaction plays a positive role in recruiting MTF2 and the PRC2 core complex to deposit H3K27me3 at genes that may have escaped Polycomb-mediated repression.

**Figure 7.**
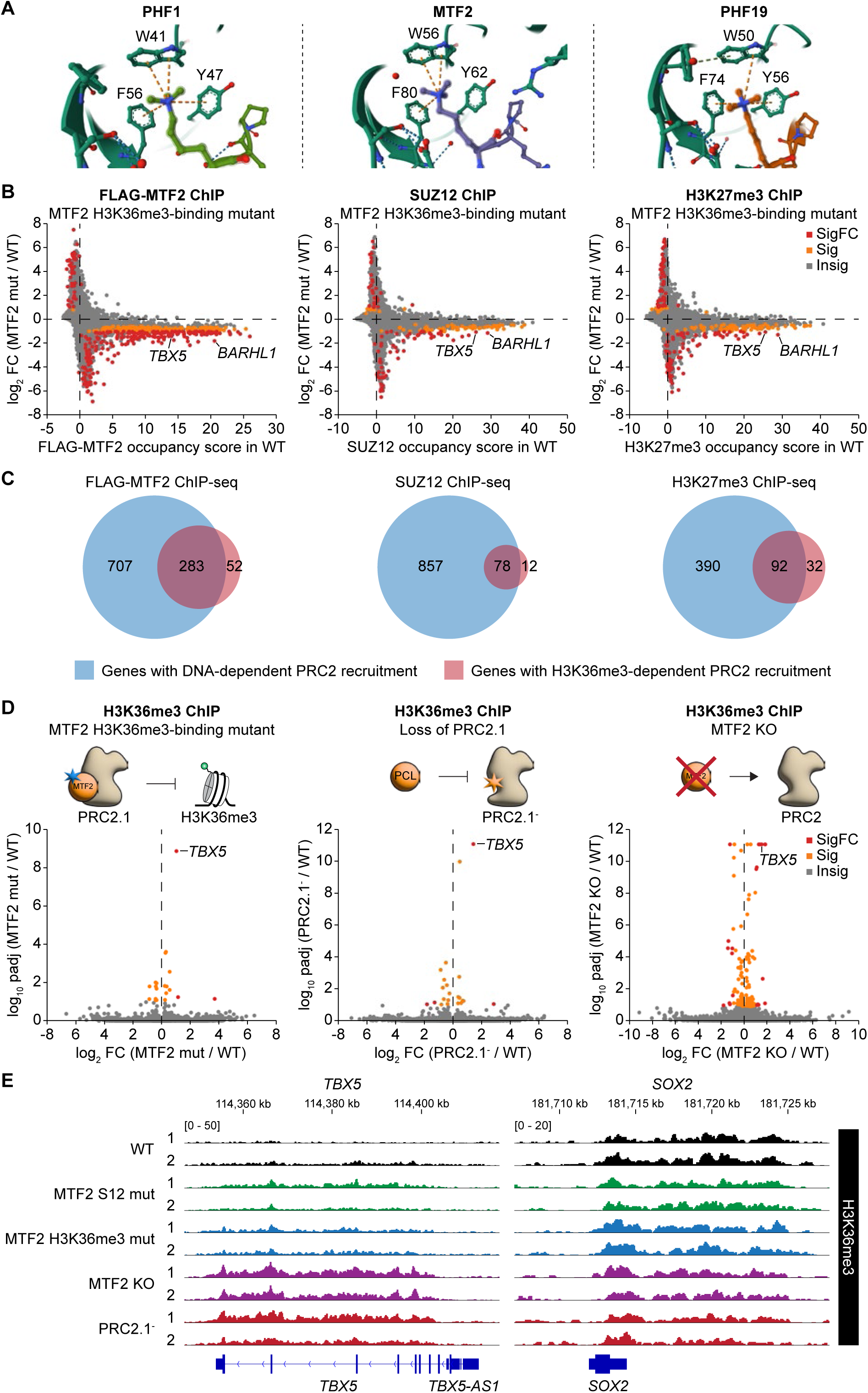
The H3K36me3-MTF2 interaction provides an alternative recruitment mechanism for PRC2 and stimulates H3K36me3 demethylation at *TBX5*. (A) Crystal structures showing the recognition of the tri-methylated lysine of H3K36me3 by the aromatic cage of the tutor domain of PHF1 (PDB: 6WAV), MTF2 (PDB: 5XFR), and PHF19 (PDB: 4BD3). The three aromatic residues are conserved in all three PCL proteins. (B) ChIP-seq gene scatter plots comparing genome-wide chromatin occupancy changes of FLAG-MTF2 (left), SUZ12 (middle), and H3K27me3 (right) between WT and H3K36me3-binding mutant (Y62A) MTF2 hiPSC lines. Two independent clones of each genotype (n = 2) were compared by empirical Wald tests for individual genes. Grey dots (Insig) indicate a multiple test-corrected (FDR) *p*-value ≥ 0.1, orange dots (Sig) an FDR-adjusted *p*-value < 0.1 and |log2FC| < 1, and red dots (SigFC) an FDR-adjusted *p*-value < 0.1 and |log2FC| ≥ 1. (C) Venn diagrams showing the number of genes with DNA- and H3K36me3-specific recruitment of MTF2 and SUZ12 and deposition of H3K27me3. The genes analyzed here represent those with an FDR-adjusted differential *p*-value (comparing WT and mutant) < 0.1 and an FDR-adjusted enrichment *p*-value (comparing pulldown and input) in the WT line < 0.1. (D) ChIP-seq volcano plots comparing the gene-specific changes of H3K36me3 levels upon mutating the aromatic cage (Y62A) of MTF2 (left), perturbing SUZ12’s interface with all three PCL proteins (middle), and depleting MTF2 (right). Grey dots (Insig) indicate a multiple test-corrected (FDR) *p*-value ≥ 0.1, orange dots (Sig) an FDR-adjusted *p*-value < 0.1 and |log2FC| < 1, and red dots (SigFC) an FDR-adjusted *p*-value < 0.1 and |log2FC| ≥ 1. (E) H3K36me3 ChIP-seq genome tracks of WT, MTF2 SUZ12-binding mutant, MTF2 H3K36me3-binding mutant, MTF2 KO, and PRC2.1^-^ at the *TBX5* and *SOX2* loci.

We then compared the genes regulated by DNA- and H3K36me3-dependent recruitment mechanisms for the MTF2-containing PRC2.1 complex. Based on the number of genes significantly changed in FLAG-MTF2, SUZ12, and H3K27me3 ChIP-seq between WT and DNA- or H3K36me3-binding mutant, DNA-mediated recruitment of MTF2 and PRC2.1 appears to play a more dominant role compared to the H3K36me3-mediated recruitment (**Figure 7C**). In addition, a majority of H3K36me3-mediated Polycomb regulation also relied on DNA-MTF2 interaction. Gene ontology analysis indicates that these overlapping genes are mostly transcription factor genes, while the genes regulated specifically by only DNA-MTF2 or H3K36me3-MTF2 interaction are involved in developmental pathways (**Figure S8A**).

We next asked how the H3K36me3-MTF2 interaction regulates H3K36me3 itself. To this end, we performed H3K36me3 ChIP-seq on the MTF2 H3K36me3-binding mutant hiPSC lines. We found that a small subset of genes exhibited altered H3K36me3 levels upon disruption of the H3K36me3-MTF2 interaction, with the cardiac transcription factor gene *TBX5* being the most significantly affected (**Figures 7D and 7E**). The increase in H3K36me3 levels on *TBX5* was consistently observed in the loss-of-PRC2.1 SUZ12 mutant, the SUZ12-binding mutant of MTF2, a second H3K36me3-binding mutant of MTF2 (F80A), as well as the MTF2 KO (**Figures 7D, 7E, and S8B**). In contrast, H3K36me3 levels on *TBX5* remained unchanged when the DNA-MTF2 interaction was perturbed (**Figure S8B**, right panel). Similarly, H3K36me3 levels on *TBX5* were not dysregulated in the SUZ12- and DNA-binding mutants or the KO of PHF1 and PHF19 (**Figures S8C and S8D**), suggesting that MTF2 is the dominant PCL protein in regulating H3K36me3 levels at the *TBX5* locus.

Transcriptomic analysis of the MTF2 H3K36me3-binding mutant revealed even larger gene expression changes than observed in the MTF2 KO (305 versus 100 dysregulated genes) (**Figures S8E and S8F**), suggesting that lacking the H3K36me3 recognition while preserving PRC2 association is more detrimental to the cell than complete depletion of MTF2. Downregulated genes were enriched in prostate gland and urogenital development, while upregulated genes were enriched in cellular respiration pathways (**Figure S8G**). All these results support a novel PRC2-dependent role of MTF2 in the gene-specific removal of H3K36me3, potentially by recruiting an H3K36me3 demethylase, a function that has so far only been proposed for PHF19 among the three PCL proteins.^19^

### Loss-of-function of MTF2 accelerates cardiac differentiation

The important role of MTF2 in the epigenetic repression of the *NKX2-5* and *TBX5* genes implicates an involvement of MTF2 in early cardiac lineage commitment and differentiation. To test this hypothesis, we induced the cardiac differentiation of the MTF2 WT, KO, and all separation-of-function mutant lines. We found that in comparison to the WT line, the MTF2 KO line exhibited a much-accelerated spontaneous contraction phenotype, followed by the separation-of-function mutant lines (**Figures 8A and 8B**). Flow cytometry analysis on day 15 of the differentiation time course, when all lines were spontaneously contracting, showed a significantly higher percentage of cTnT-expressing cells for all MTF2 loss-of-function lines compared to the WT (**Figures 8C and S9A**). This difference could already be observed on day 8 but was less pronounced than on day 15 (**Figures S9A and S9B**). Of note, differentiation of the PHF1 KO lines similarly yielded more cTnT-expressing cells in comparison to the WT, while no significant difference could be observed between PHF19 WT and KO (**Figure S9C**).

**Figure 8.**
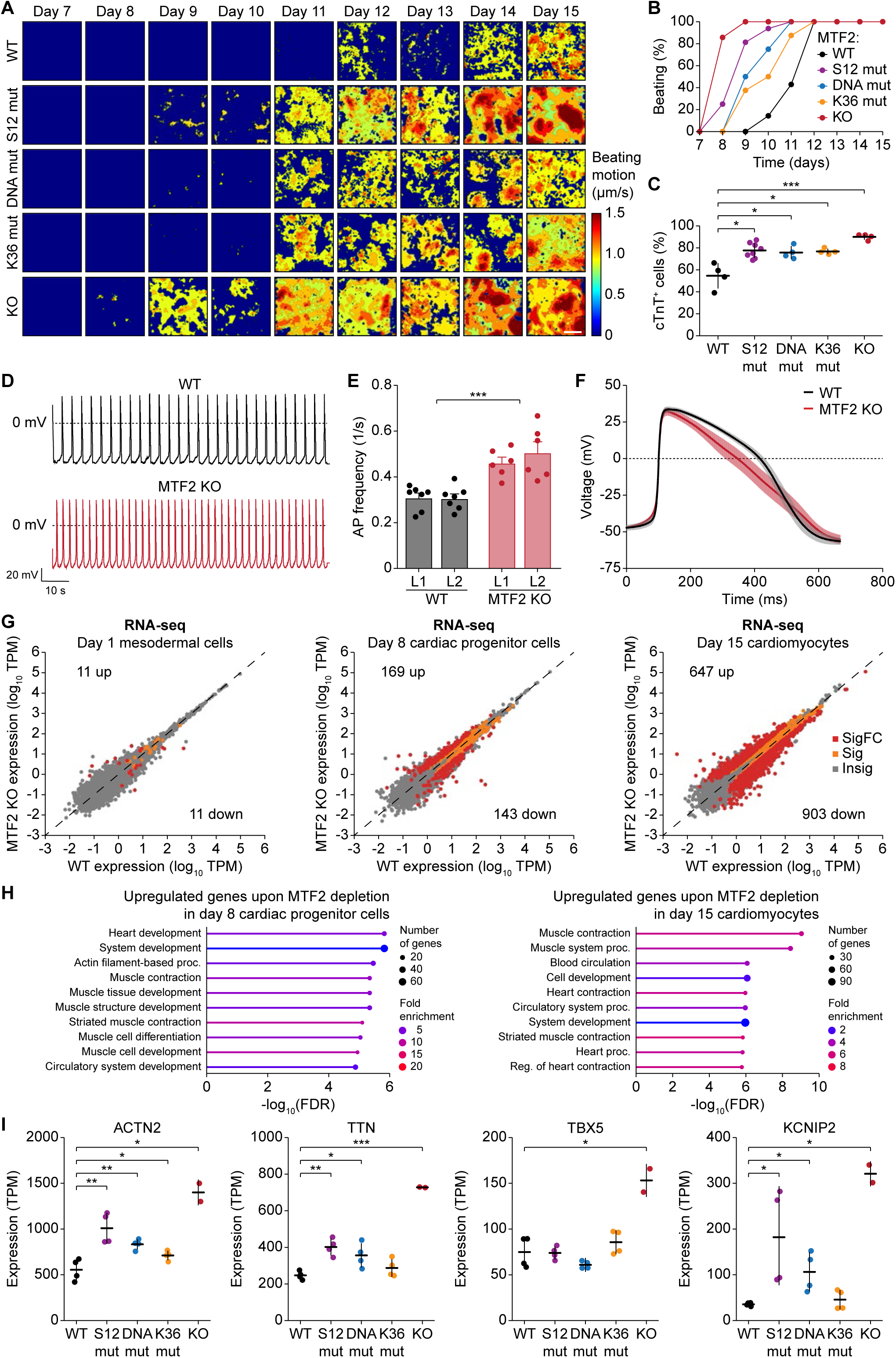
Loss-of-function of MTF2 accelerates cardiac differentiation. (A) Heatmaps showing the time-averaged magnitude of spontaneous contractions of WT and mutant MTF2 cardiomyocytes along the differentiation process. Each heatmap is representative of eight (all but one) or 16 (SUZ12-binding mutant) independent recordings. (B) Percentage of cell culture wells with spontaneous contractions of WT and mutant MTF2 cardiomyocytes over time. Data are from each genotype cultured in multiple wells (n = 14 for WT, n = 16 for SUZ12-binding mutant, n = 8 for DNA-binding mutant, n = 8 for H3K36me3-binding mutant, and n = 14 for MTF2 KO). (C) Percentage of WT and mutant MTF2 cells expressing cTnT after 15 days of differentiation as determined by flow cytometry. Data are shown as mean and standard deviation of each genotype (n = 4 for WT, n = 8 for SUZ12-binding mutant, n = 4 for DNA-binding mutant, n = 4 for H3K36me3- binding mutant, and n = 4 for MTF2 KO). * *P* < 0.05, *** *P* < 0.001, two-tailed Student’s *t*-test. (**D-F**) Electrophysiological analysis of WT and MTF2 KO day 24 cardiomyocytes as recorded by patch clamp. Representative action potential (AP) traces (D), bar plots of average AP frequency (E), and average single AP trace of all measured cells over time (F) of spontaneous contractions are shown. Data in (E and F) are shown as mean and standard error of the mean of seven cell measurements. *** *P* < 0.001, two-way ANOVA. (G) RNA-seq scatter plots showing gene expression changes upon loss of MTF2 at day 1, 8, and 15 of the cardiac differentiation process. Two independent clones of each genotype cultured independently twice (n = 4) were compared for statistical significance (two-sided Wald test). Grey dots (Insig) indicate a multiple test-corrected (FDR) *p*-value ≥ 0.05, orange dots (Sig) an FDR-adjusted *p*-value < 0.05 and |log2FC| < 1, and red dots (SigFC) an FDR-adjusted *p*-value < 0.05 and |log2FC| ≥ 1. Number of up- and downregulated genes (SigFC, red dots) are indicated. (H) Gene ontology analysis of the upregulated genes upon depletion of MTF2 in day 8 cardiac progenitor cells and day 15 cardiomyocytes. (I) Gene expression of *ACTN2*, *TTN*, *TBX5*, and *KCNIP2* in day 15 cardiomyocytes as determined by RNA-seq. Data are shown as mean and standard deviation of two (MTF2 KO) or four (WT and variant MTF2) biological replicates each. * *P* < 0.05, ** *P* < 0.01, *** *P* < 0.001, two-tailed Student’s *t*-test.

We then used metabolic selection with lactate to enrich the cardiomyocyte population. Immunofluorescence staining revealed that the MTF2 KO cardiomyocytes exhibited significantly lower H3K27me3 levels compared to the WT (**Figures S9D and S9E**), similar to what we observed in the undifferentiated hiPSCs (**Figures 1C, S1E, and S1F**), suggesting an important role of MTF2 in H3K27me3 maintenance in cardiomyocytes. Interestingly, the PHF19 KO cardiomyocytes also exhibited lower H3K27me3 levels, inferring a positive role of PHF19 in H3K27me3 deposition in cardiomyocytes (**Figures S9D and S9E**). Importantly, patch clamp recordings of day 24 cardiomyocytes showed that the electrophysical properties of the MTF2 KO line phenocopied those of the loss-of-PRC2.1 line. Depletion of MTF2 led to significantly higher action potential frequency (**Figures 8D and 8E**) and faster depolarization (**Figure 8F**), with unperturbed action potential peak amplitude and duration (**Figures S9F and S9G**).

We lastly asked what transcriptomic changes ensued during the cardiac differentiation time course. RNA-seq analysis comparing MTF2 WT and KO lines collected on day 0, 1, 8, and 15 of cardiac differentiation suggests more pronounced global gene expression changes in later stages of the differentiation process (**Figure 8G**). This observation is consistent with the perturbation of epigenetic repression of developmental genes upon MTF2 depletion in the undifferentiated state (**Figures 4H and 4I**). Gene ontology analysis of the upregulated genes in day 8 cardiac progenitor cells and day 15 cardiomyocytes suggests an increased expression of cardiac genes (**Figure 8H**).

For example, most MTF2 mutant and all MTF2 KO lines showed an elevated expression of *ACTN2*, *TTN*, *TBX5*, and *KCNIP2* on day 15 (**Figure 8I**). Interestingly, the three separation-of-function mutants of MTF2 exhibited unique gene expression profiles (**Figure S9H**), implicating specified roles of these macromolecular interactions in gene expression during cell differentiation. In summary, we demonstrate that MTF2 is the most important PCL protein in cardiomyocyte differentiation, as MTF2 loss-of-function accelerates cardiac differentiation and increases action potential frequency by upregulating cardiac and ion channel genes.

## Discussion

The molecular mechanisms to specifically tune epigenetic repression during cell differentiation remain one of the unsolved conundrums in biology. Recent studies using gene or protein depletion strategies have identified the accessory proteins of PRC2 as the key regulators of Polycomb-mediated epigenetic repression.^4, 12, 14, 22, 33^ However, PRC2 accessory proteins are involved in versatile macromolecular interactions, and thus complete depletion may limit an understanding of their molecular mechanism. In this study, we performed “surgical operations” using a series of separation-of-function mutations to precisely modify a minimal number of amino acid residues. This approach allowed us to disrupt protein-protein, protein-DNA, and protein-histone modification interactions that mediate the recruitment process of PRC2.1, PRC2.2, and the three mutually exclusive PCL protein-containing PRC2.1 complexes. Contrary to previous conclusions on the redundancy and largely overlapping roles of these subcomplexes, our study demonstrates the distinct specificity and role of each subcomplex in epigenetic repression.

### Distinct versus overlapping roles for PRC2.1 and PRC2.2

The PRC2.1 and PRC2.2 subcomplexes were previously proposed to function redundantly.^14, 33^ However, our study suggests gene-specific roles for PRC2.1 and PRC2.2 in depositing H3K27me3. While both recombinant complexes were biochemically more active than the PRC2 core complex, PRC2.1 and PRC2.2 specifically deposited H3K27me3 on two distinct subsets of genes (577 versus 361 genes, with only 30 overlapping genes). While PRC2.1-repressed genes were enriched for developmental transcription factor genes, PRC2.2-repressed genes were more functionally diverse. The distinct roles of PRC2.1 and PRC2.2 were further demonstrated by the distinct changes in PRC1 components and the transcriptome. Loss of PRC2.2 led to substantially larger gene expression changes compared to the loss of PRC2.1, indicating that most of the developmental genes regulated by PRC2.1 have yet to be transcriptionally activated in undifferentiated cells. The functional divergence between PRC2.1 and PRC2.2 may have evolved to enable increased regulatory complexity and robustness during development and in response to environmental cues.

### Unidirectional competition between of PRC2.1 and PRC2.2

The competition between the two sets of accessory subunits in PRC2.1 and PRC2.2 appears to be unidirectional. Loss of PRC2.2 reinforced the interaction between SUZ12 and PHF1/MTF2 and led to a gain in SUZ12 chromatin occupancy, which can be explained by the higher chromatin affinity of PRC2.1 compared to PRC2.2.^13^ In contrast, loss of PRC2.1 did not lead to any significant change in PRC2.2 complex composition or chromatin occupancy. This unidirectional phenomenon could be due to the stoichiometry of the accessory proteins in PRC2.1 and PRC2.2. A reasonable hypothesis is that there are free/unbound PCL proteins (especially PHF1) available to bind to the newly exposed PRC2 core complex when PRC2.2 is disrupted. In contrast, no unbound JARID2 or AEBP2 proteins are available to form more PRC2.2 complexes upon disruption of PRC2.1.

### Distinct and overlapping roles of the three PCL proteins

We also identified the distinct roles for the three PCL proteins for the first time. While it was previously thought that the PCL proteins have redundant functions in regulating PRC2,^33^ we showed in this study that the three PCL proteins employ distinct regulatory mechanisms and specificities. The results were consistent across a series of approaches (depletion, mutation of PCL protein-SUZ12 interaction, and mutation of PCL protein-DNA interaction). Especially PHF19, the most lowly expressed PCL protein in undifferentiated stem cells, was shown to have an antagonistic role by competing for target genes with the MTF2-PRC2.1 subcomplex. This is potentially a self-regulatory mechanism to prevent hyperactivity of the enzyme complex. The PHF19-PRC2.1 complex may interact with additional inhibitor proteins to keep the enzyme in a poised and chromatin-unbound state. Further investigation is needed to identify such proteins.

### Recruitment of PRC2.1 by DNA versus H3K36me3

We found that both DNA and H3K36me3 contribute to the recruitment of PRC2.1 in human stem cells. It remains to be uncovered how these two recruitment mechanisms crosstalk and work synergistically to establish epigenetic repression. We also identified a gene-specific role of MTF2 in maintaining low H3K36me3 levels on the *TBX5* gene in hiPSCs, which is mediated through the interaction between H3K36me3 and the Tudor domain of MTF2. Interestingly, the PHF19-H3K36me3 interaction has been reported to recruit an H3K36 demethylase to enhance epigenetic repression.^19^ MTF2 has a similar function to maintain low H3K36me3 levels at the *TBX5* gene to prevent premature activation before cardiac lineage commitment.

### Polycomb-mediated epigenetic regulation of cardiac differentiation

PRC2 is essential for heart development and functions, but previous studies were heavily focused on the subunits in the core complex.^31, 42, 43^ Beyond the core complex, we show here that the PRC2 subcomplexes play important roles in regulating stem cell cardiac differentiation by either promoting (PRC2.2) or repressing (PRC2.1) cardiac lineage commitment. On one hand, perturbation of PRC2.1 – either by depletion of MTF2 or disruption of SUZ12-PCL protein interaction, SUZ12-MTF2 interaction, MTF2-DNA interaction, or MTF2-H3K36me3 interaction – led to the stimulation and acceleration of cardiac differentiation. On the other hand, perturbation of PRC2.2 – by disruption of the SUZ12-AEBP2/JARID2 interaction – negatively impacted cardiac differentiation. These results suggest that PRC2.1, particularly the MTF2-PRC2.1 complex, represses cardiac lineage commitment, while PRC2.2 is required for cardiac differentiation. The negative role of PRC2.1 in cardiac differentiation is potentially mediated through the epigenetic repression of key cardiac transcription factor genes such as *NKX2-5* and *TBX5* as well as important ion channel genes, including *KCNQ1*, *KCNIP2*, and *HCN2.* These observations are consistent with the identified roles of PRC2 in regulating cardiac transcription factors^31, 42, 43^ and ion channels^44^ during differentiation and normal heart function. In contrast, the requirement of PRC2.2 in cardiac differentiation could be mediated by the specific repression of genes that inhibit cardiac lineage commitment such as BMP antagonist gene *CER1* and ectopic lineage transcription factor genes *FOXA2*, *LHX5*, and *OLIG3* (**Table S3**). This suggests that the PRC2 subcomplexes provide a balance in lineage commitment that could facilitate the desired cellular output in development and regeneration after injury. This study opens the door to future investigations of PRC2 accessory proteins in the heart, especially since JARID2,^45–47^ PHF1,^48^ MTF2,^4, 49^ and PHF19^50^ have been implicated in cardiac development and function. Furthermore, the key macromolecular interactions investigated in this study could also impact Polycomb-mediated epigenetic outcome in other tissues beyond the heart, especially since a variety of lineage-specific transcription factor genes were found to be dysregulated upon loss of PRC2.1 or PRC2.2 (**Table S2-3**). Future differentiation experiments or organoid models will be valuable to further identify the roles of the PRC2 subcomplexes in various developmental contexts.

The two transcription factor genes *NKX2-5* and *TBX5*, which are master regulators of cardiac development, were found to be especially sensitive to the perturbation of the PRC2.1 subcomplexes. A potential reason why they frequently appeared as the most affected genes regarding PRC2 occupancy or H3K27me3 levels is the high CpG content in their gene body and genetic neighborhood, making them high affinity PRC2.1 targets. Developing therapies that target PRC2-mediated epigenetic regulation of *NKX2-5* and *TBX5* may enable the treatment of congenital heart diseases, where the transcription of these two genes is frequently dysregulated.^51, 52^

### Separation-of-function residues are frequently mutated in cancer

The amino acid residues that we targeted in our separation-of-function studies are frequently mutated in cancer. Mutations of all three SUZ12-interacting residues of MTF2 (R574G/W, L584P and W587L) have been found in various types of carcinoma.^53–55^ Similarly, mutations of the DNA-binding and H3K36me3-binding residues of MTF2 (K338Rfs*51 and F80C) have been identified in different cancers.^55^ PCL-interacting residues (338-353) of SUZ12 are also frequently mutated in cancers, including L338V/H, D339G, G340V/R, R342W, P345L, E347K/D, F349L, G352R and P353H.^56–62^ Among the four AEBP2/JARID2-interacting residues of SUZ12 (F99, R103, L105 and I106), R103 seems to be particularly prone to mutations in cancer (R103Q/P/W), and it has been shown that either of the R103Q or R103P mutation is sufficient to perturb the balance between PRC2.1 and PRC2.2.^13^ Future investigation will be needed to identify the roles of these disease mutations in PRC2 subcomplex formation and epigenetic repression.

### Concluding remarks

Our study employed separation-of-function mutants to “surgically” perturb macromolecular interactions, which is extremely suitable for the mechanistic study of the regulation of epigenetic repression in stem cells and differentiation. These findings not only uncovered the distinct roles of PRC2 subcomplexes and how epigenetic specificity is precisely controlled, but also set a new paradigm for the mechanistic study of complex epigenetic regulation. Our mechanistic study sets the stage for future drug development to perturb these macromolecular interfaces and suppress Polycomb-mediated repression in a gene-specific manner, as well as potential epigenome editing to correct epigenetic defects in human diseases.

### Limitations of the study

Our genome editing approach enabled us to introduce a WT or mutant cDNA copy at the endogenous gene locus. This approach may potentially alter the regulation of mRNA isoforms. It is possible that alternative isoforms could exist at different differentiation stages and could increase the functional complexity and diversity of PRC2.

We used the common hiPSC line WTC-11, which is derived from a male individual, to generate all our CRISPR-edited cell lines. It cannot be ruled out that hiPSCs from a different donor could behave differently, or that female cells could show a different phenotype, although a sex-specific function of PRC2 outside of X-inactivation has not been reported so far.

We characterized the functional importance of various PRC2 components and macromolecular interactions in cardiac differentiation. This does not preclude an important role for PRC2.1 and PRC2.2 in other cell types and lineages. For instance, the PRC2.1 subcomplexes could show very different behaviors based on the expression pattern of the individual PCL proteins.

## STAR★Methods

### Key resources table

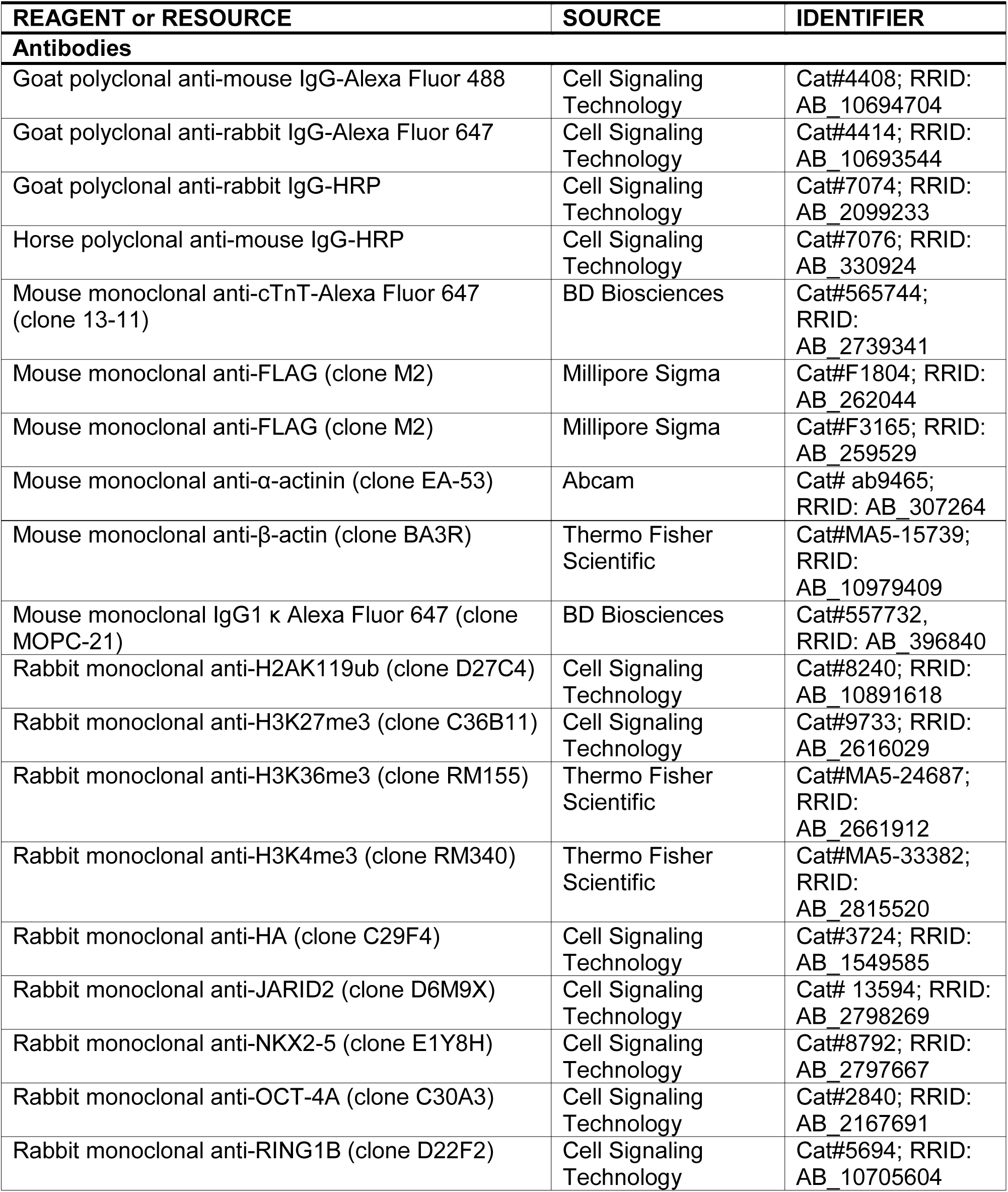
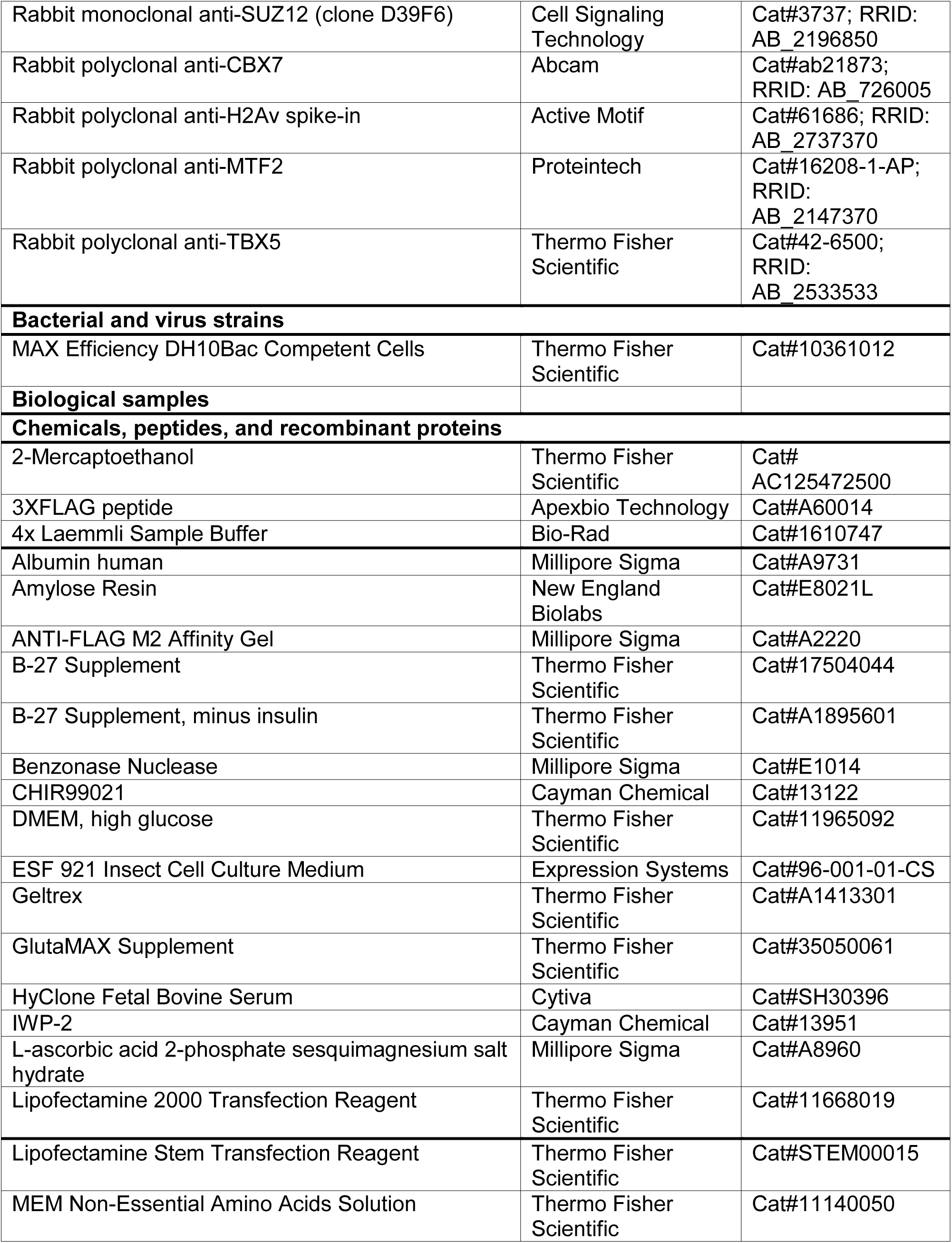
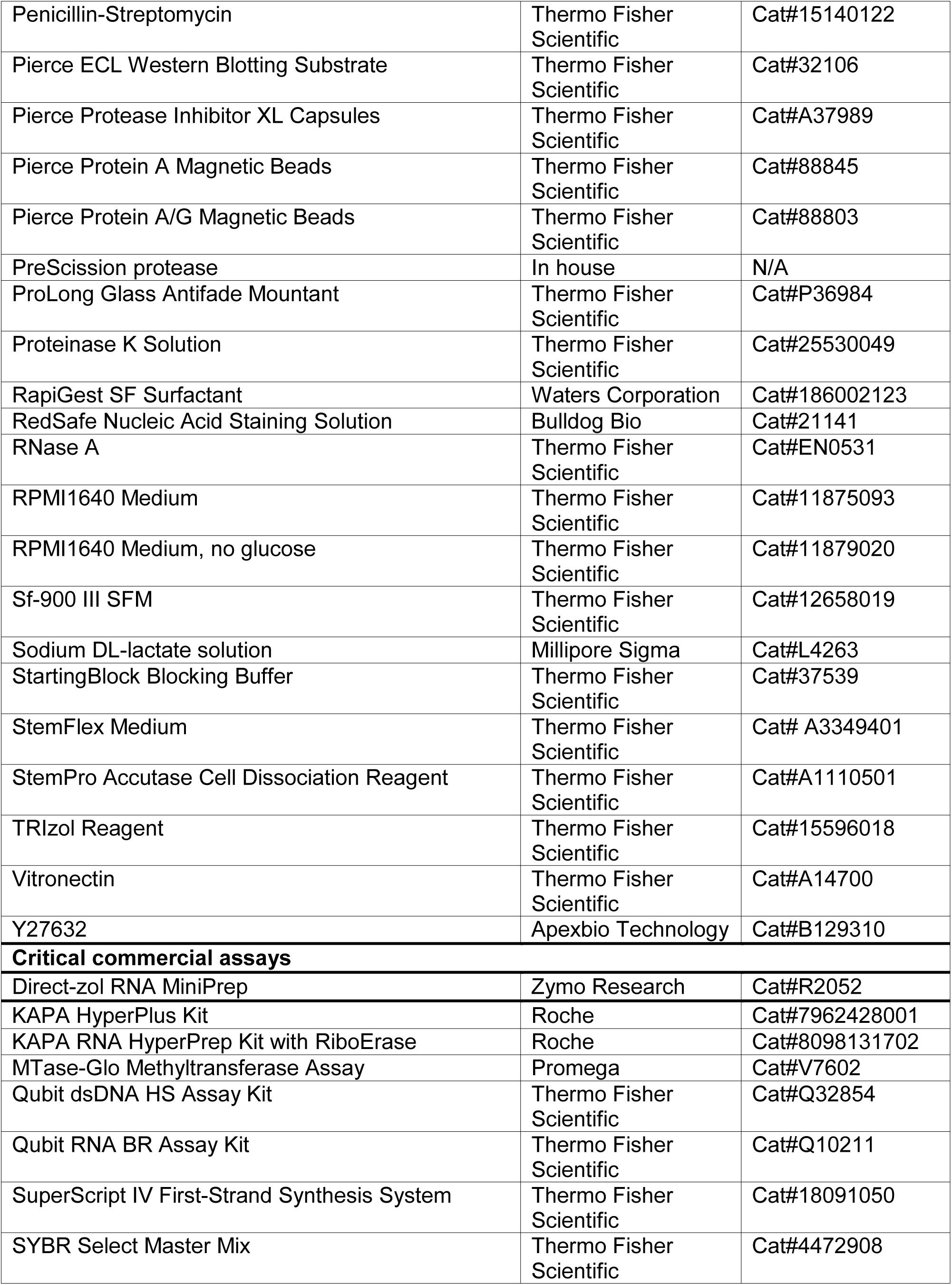
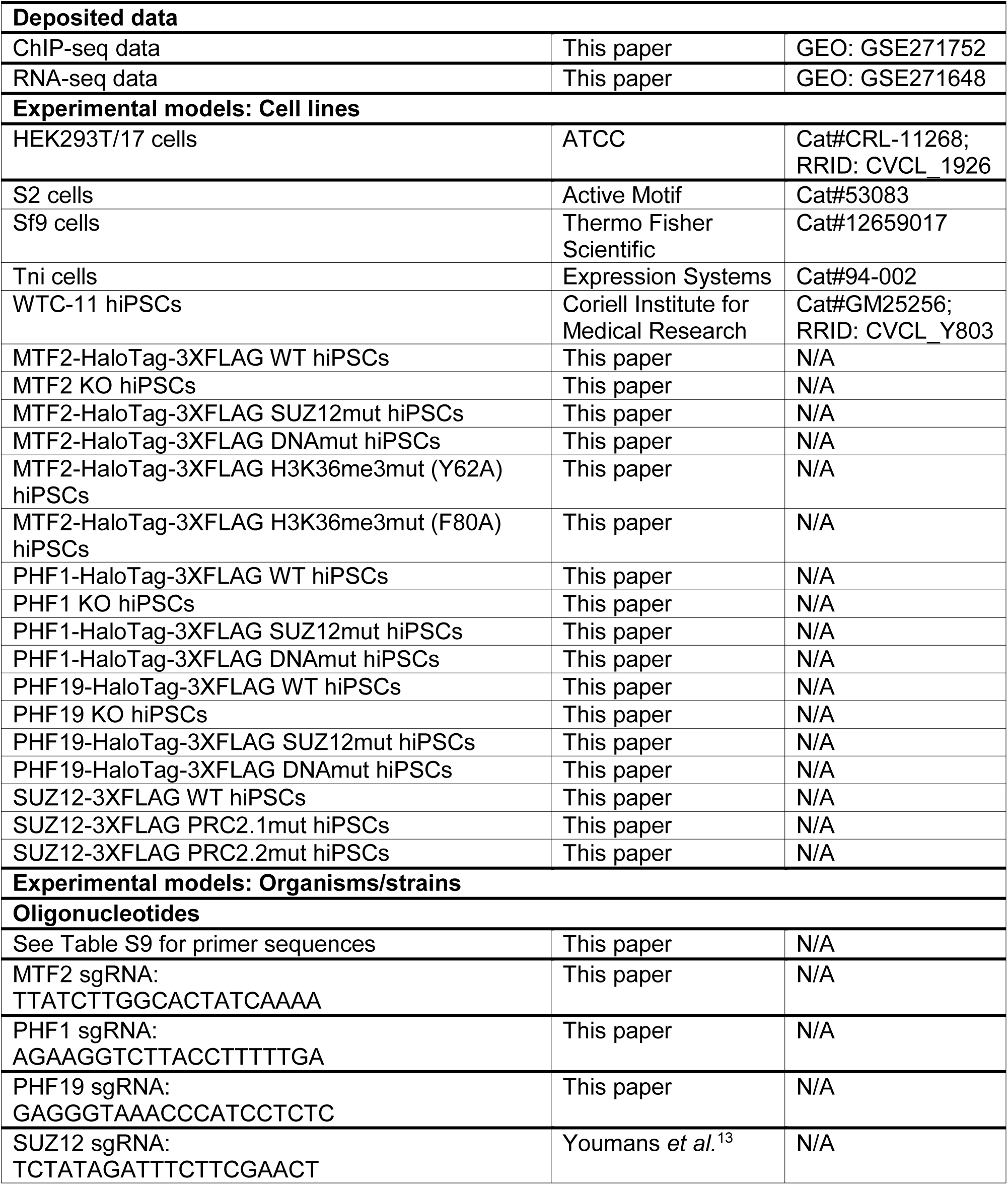
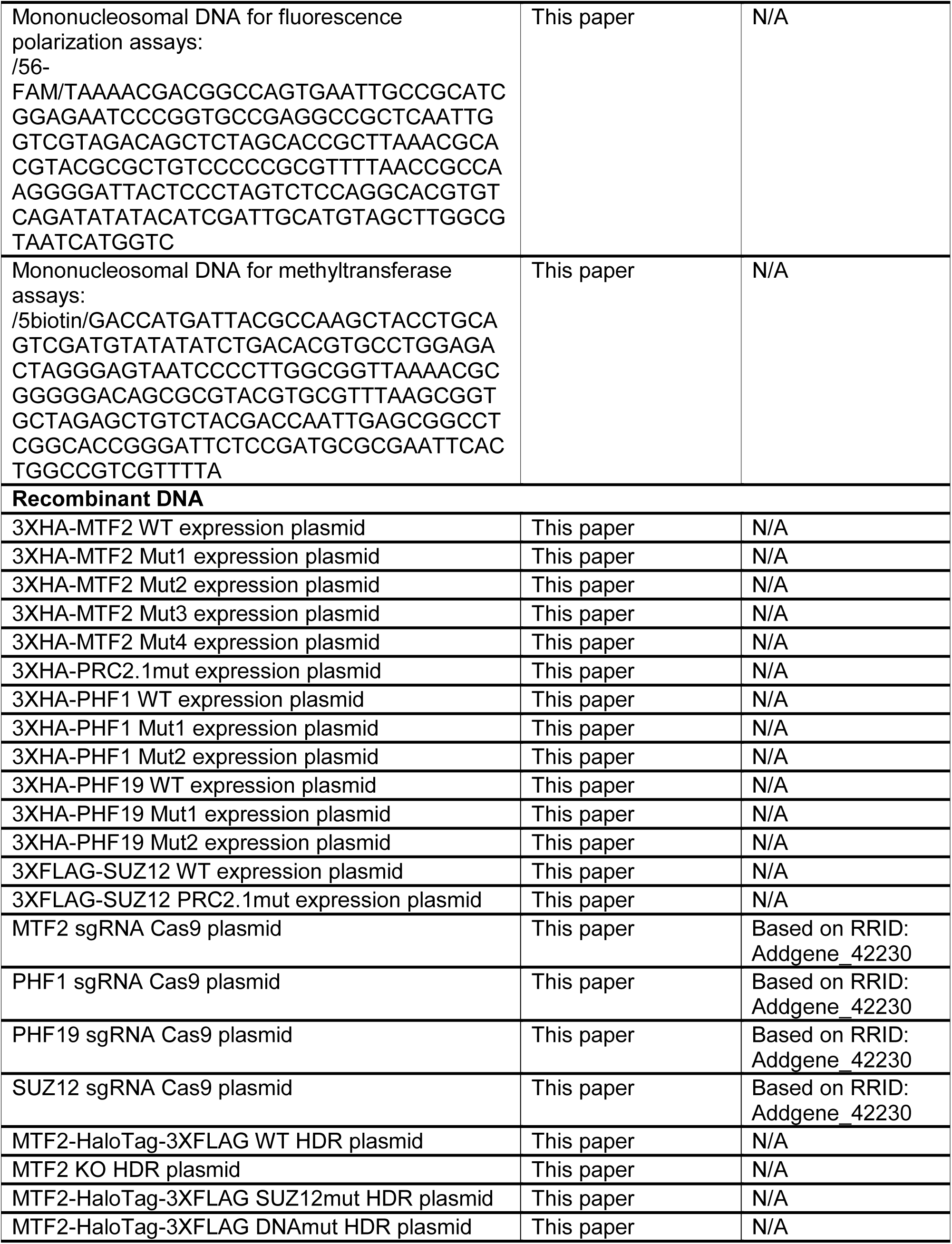
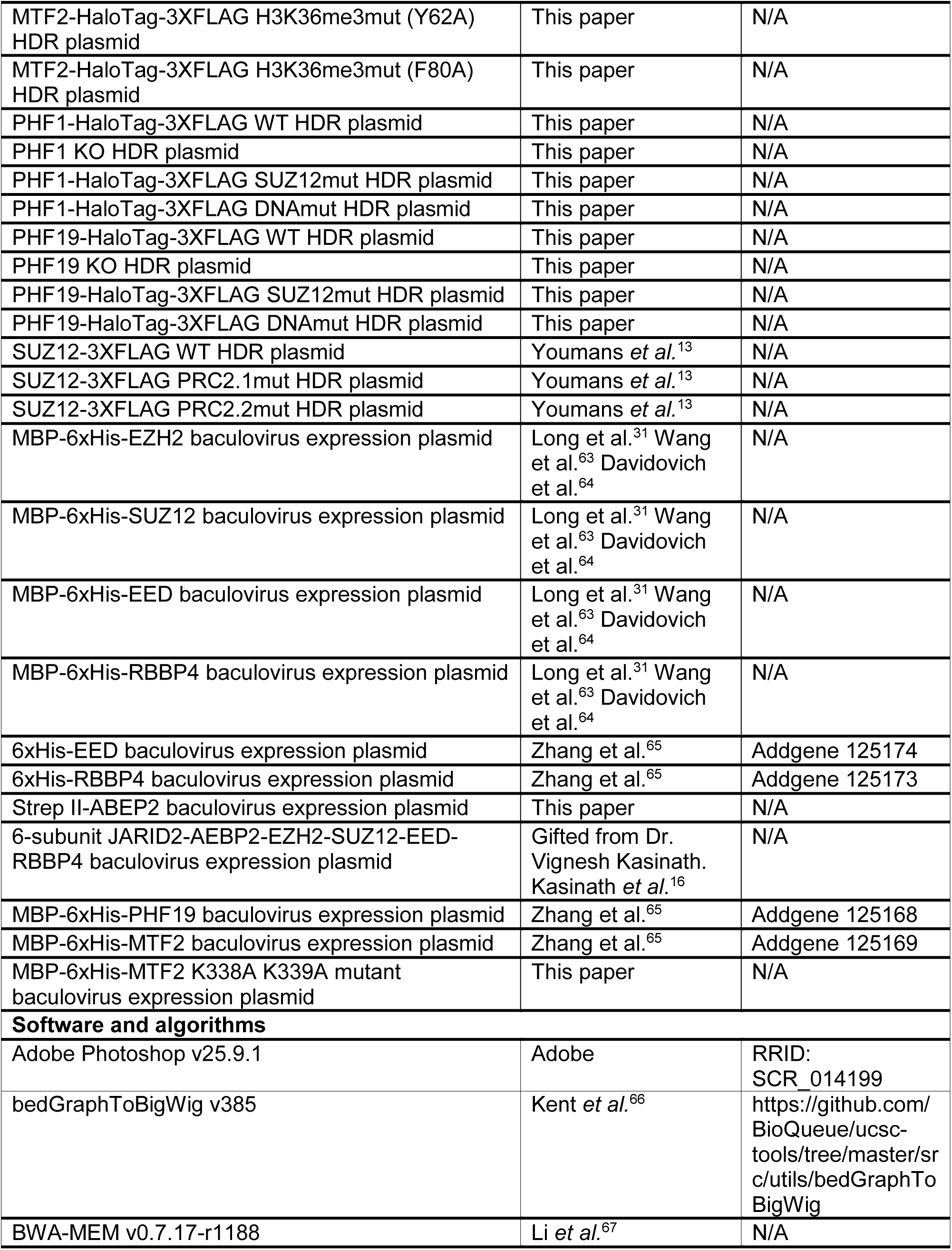
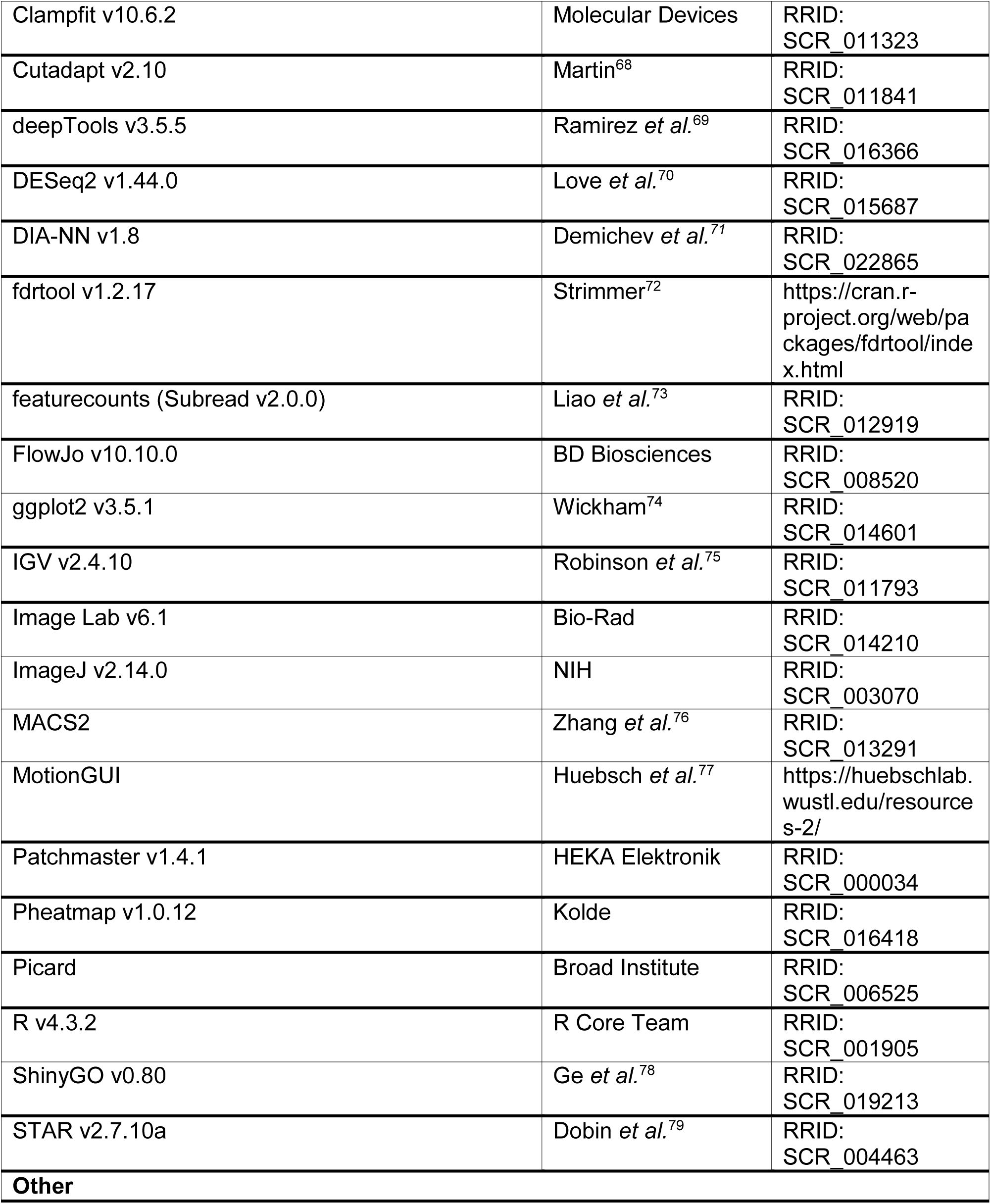

### Resource availability

Further information and requests for resources and reagents should be directed to and will be fulfilled by the lead contact, Yicheng Long.

### Materials availability

All cell lines and plasmids generated in this study are available from the lead contact with a completed material transfer agreement.

### Data and code availability

ChIP-seq and RNA-seq data have been deposited at GEO under accession number GSE271752 and GSE271648 and are publicly available as of the date of publication. Original western blot images are shown in Data S1.

The paper does not report original code.

Any additional information required to reanalyze the data reported in this paper is available from the lead contact upon request.

### Experimental model and study participant details

#### hiPSC culture

Male hiPSCs were maintained on vitronectin in StemFlex medium supplemented with 25 U/mL penicillin/streptomycin at 37°C and 5% CO_2_. Cells were not authenticated. Cells were routinely tested for mycoplasma contamination by PCR.

#### HEK293T cell culture

Female HEK293T cells were maintained in DMEM supplemented with 10% fetal bovine serum, 1X GlutaMAX supplement, 1X non-essential amino acids, 100 U/mL penicillin/streptomycin, 100 μM 2-mercaptoethanol at 37°C and 5% CO_2_. Cells were not authenticated. Cells were routinely tested for mycoplasma contamination by PCR.

### Method details

#### CRISPR genome editing

The CRISPR plasmid encoding Cas9 and guide RNA was generated by inserting the guide RNA sequence targeting the junction between exon 2 and intron 2 of the gene to be edited into pX330. The donor plasmids carrying either the WT or mutated complementary DNA were constructed by assembling the following fragments into a previously described donor plasmid:^80^ left homology arm, target gene cDNA, tag, 3′ UTR, 3x SV40 polyadenylation sites, 1X bGH polyadenylation site, hPGK promoter, puromycin or blasticidin resistance ORF, 1X bGH polyadenylation site, and right homology arm. KOs were generated with similar donor plasmids but lacking the cDNA, tag, and 3’UTR portion. A total of 2.5 µg of equal amounts of CRISPR plasmid and two instances of donor plasmids carrying either the puromycin or blasticidin resistance ORF were transfected into WTC-11 hiPSCs in a 6-well plate using Lipofectamine Stem Transfection Reagent following the manufacturer’s instructions. After two to three days, cells were passaged to a 10-cm plate coated with Geltrex. Once cells reached 60-80% confluency, 1 μg/ml puromycin was added to the culture for four days. Cells were then selected in the presence of both 1 μg/ml puromycin and 5 μg/ml blasticidin for one week. At least 24 surviving colonies were manually picked, transferred to 24-well plates, and expanded. Successfully genome-edited colonies were verified by PCR, Sanger sequencing, and western blotting. Two or three independent clones of genome-edited hiPSC lines were used in subsequent experiments.

#### Western blotting

Cells were collected by Accutase treatment, washed with cold PBS, and incubated in lysis buffer (25 mM Tris pH 7.5, 150 mM NaCl, 2.5 mM MgCl2, 1% Igepal, 5% glycerol, 2 mM TCEP, 1X protease inhibitor cocktail, 1 µL/mL benzonase) for 10 min on ice. Protein extracts were collected as supernatant after centrifugation at 16,000 x g for 10 min at 4°C, incubated in 1X Laemmli buffer for 10 min at 95°C, resolved on a 4-15% Mini-PROTEAN TGX Stain-Free Protein Gel (Bio-Rad, Cat#4568084), and transferred to a nitrocellulose membrane via wet transfer. Membranes were blocked in StartingBlock Blocking Buffer and probed with anti-β-actin, anti-FLAG, anti-HA, anti-H3K27me3, anti-SUZ12, or anti-MTF2 at a dilution of 1:1000 in blocking buffer overnight at 4°C or for 2 h at room temperature. The membrane was washed in PBS-T (0.1% Tween-20 in PBS), incubated with an anti-mouse or anti-rabbit HRP-coupled secondary antibody in PBS-T for 1 h, and washed again in PBS-T. Proteins were detected with the Pierce ECL Western Blotting Substrate and acquired using a ChemiDoc Imaging System (Bio-Rad). Data were analyzed using Image Lab.

#### Immunofluorescence staining

Cells grown on cover slips were fixed with 4% formaldehyde in PBS for 10 min and then permeabilized in extraction buffer (0.5% Triton X-100, 20 mM HEPES-KOH pH 7.5, 50 mM NaCl, 3 mM MgCl2, 300 mM sucrose) for 10 min. Cover slips were then washed with PBS + 0.1% Triton X-100 twice and blocked in ABDIL buffer (3% BSA and 0.1% Triton X-100 in PBS) for 30 min. Samples were subsequently incubated in anti-α-actinin (1:200), anti-cTnT-Alexa Fluor 647 (1:200), anti-FLAG (1:400), anti-H3K27me3 (1:200), anti-NKX2-5 (1:400), anti-OCT-4 (1:200), or anti-TBX5 (1:200) primary antibody diluted in ABDIL buffer for a 1 h, washed three times in PBS, and incubated in anti-mouse Alexa Fluor 488 or anti-rabbit Alexa Fluor 647-conjugated secondary antibody diluted 1:500 in ABDIL buffer containing 10 μg/mL Hoechst 33342 (Thermo Fisher, H3570) for 30 min. After another three PBS washes, cover slips were mounted onto a glass slide using ProLong Glass Antifade Mountant. Images were acquired as Z-stacks of 200-nm step size on a Leica DMi8 microscope with a Hamamatsu ORCA-Fusion BT digital camera and were equally processed using ImageJ/Fiji and Photoshop.

#### Immunoprecipitation followed by mass spectrometry

hiPSCs at 70-80% confluency were collected by Accutase treatment, washed with PBS, and extracted with 1 mL IP-MS lysis buffer (50 mM Tris pH 8, 100 mM KCl, 5 mM MgCl2, 10 % glycerol, 0.5 % NP-40, 1X protease inhibitor cocktail, 1 µL/mL benzonase) on ice for 10 min. Protein extracts were collected as supernatant after centrifugation at 16,000 x g for 10 min at 4°C. Pierce Protein A Magnetic Beads crosslinked to anti-SUZ12 were added to the lysate and incubated at 4°C for 30 min. Beads were washed three times with IP-MS lysis buffer without glycerol and twice with IP-MS wash buffer (50 mM Tris pH 8, 0.5 % NP-40) at 4°C. Samples were eluted by incubating the beads in IP-MS elution buffer (50 mM Tris pH 8, 0.5 % SDS) at 50°C for 15 min and magnetically removing the beads. Quality of the IP samples was confirmed by silver staining and western blotting.

For mass spectrometry, protein samples were acetone precipitated and resuspended in 0.1% RapiGest SF Surfactant and 25 mM ammonium bicarbonate. The samples were then reduced with DTT, alkylated with iodoacetamide, and digested with trypsin overnight at 37°C. The digests were desalted by C18-StageTip columns. The digests were analyzed using an EASY-nLC 1200 (Thermo Fisher Scientific) coupled online to an Orbitrap Fusion Lumos mass spectrometer (Thermo Fisher Scientific). Buffer A (0.1% FA in water) and buffer B (0.1% FA in 80% ACN) were used as mobile phases for gradient separation. A 75 µm x 15 cm chromatography column (ReproSil-Pur C18-AQ, 3 µm, Dr. Maisch HPLC) was packed in-house for peptide separation. Peptides were separated with a gradient of 5-40% buffer B over 30 min and 40%-100% buffer B over 10 min at a flow rate of 400 nL/min. The mass spectrometer was operated in a data-independent acquisition (DIA) mode. MS1 scans were collected in the Orbitrap mass analyzer over 350-1400 m/z at a resolution of 120K. The instrument was set to select precursors in 45 x 14 m/z wide windows with 1 m/z overlap from 350-975 m/z for HCD fragmentation. The MS/MS scans were collected in the orbitrap at 15K resolution. Data were searched against the human Uniprot database (8/7/2021) using DIA-NN and filtered for 1% false discovery rate for both protein and peptide identifications.

#### ChIP-seq

hiPSCs at 70-80% confluency were collected by Accutase treatment and crosslinked with 1% formaldehyde in PBS at room temperature for 10 min. The crosslinking reaction was quenched by adding one volume of 1.25 M glycine to 10 volumes of PBS for 2 min. Cells were washed in ice-cold PBS and cell pellets stored at -80°C. Aliquots of 10 million cells were lysed in 1 mL ChIP lysis buffer (800 mM NaCl, 25 mM Tris pH 7.5, 5 mM EDTA, 1% Triton X-100, 0.1% SDS, 0.5% sodium deoxycholate, 1X protease inhibitor cocktail) on ice for 30 min. Lysates were sonicated in MilliTubes (Covaris, Cat#520135) with the Covaris S220 focused-ultrasonicator at 4°C for 10 min at a PIP of 140 W, a duty factor of 5%, and 200 cycles/burst. Lysate was cleared by centrifugation at 16,000xg for 10 min at 4°C, and supernatant was collected. For each IP, 1 mL chromatin dilution buffer (25 mM Tris pH 7.5, 5 mM EDTA, 1% Triton X-100, 0.1% SDS, 1X protease inhibitor cocktail) together with 20 ng spike-in chromatin from *Drosophila melanogaster* S2 cells was added to the cleared lysate from 2.5-5.0 million cells. Chromatin samples were incubated with 2-4 μg of anti-CBX7 (4 μL), anti-FLAG (2 μL), anti-H2AK119ub (5 μL), anti-H3K27me3 (6 μL), anti-H3K36me3 (3 μL), anti-H3K4me3 (4 μL), anti-JARID2 (4 μL), anti-MTF2 (5 μL), anti-RING1B (7 μL), or anti-SUZ12 (4 μL) antibody at 4°C overnight. A spike-in antibody against the Drosophila-specific histone variant H2Av was added to all chromatin samples. Then, 25 μl of washed Pierce Protein A/G Magnetic Beads were added to the IP solution to incubate at room temperature for 2 h. Beads were washed twice with 1 mL of LSB buffer (20 mM Tris pH 8.0, 2 mM EDTA, 150 mM NaCl, 0.1% SDS, 1% Triton X-100), HSB buffer (20 mM Tris pH 8.0, 2 mM EDTA, 500 mM NaCl, 0.1% SDS, 1% Triton X-100), LiCl buffer (10 mM Tris pH 8.0, 1 mM EDTA, 250 mM LiCl, 1% sodium deoxycholate, 1% Igepal), and TE buffer (10 mM Tris pH 8.0, 1 mM EDTA). Chromatin was then eluted by incubating the beads with 120 μl of elution buffer (100 mM sodium bicarbonate, 1% SDS) for 20 min at room temperature. Formaldehyde crosslinking was reversed by adding 5 μL of 5 M NaCl and incubating at 65°C for 3 h. Protein and RNA were removed by adding 14 μL of 1M Tris pH 8, 3 μL of 500 mM EDTA, 3 μL of proteinase K, and 1 μL of 1 mg/mL RNase A and incubating at 37°C for 1 h. DNA was extracted by phenol-chloroform-isoamyl extraction followed by ethanol precipitation. DNA samples were resuspended in 30 μl of 10 mM Tris pH 8.0 and quantified using a Qubit fluorometer (Invitrogen). Between 2 ng and 100 ng of each sample was used for library preparation using the KAPA HyperPlus Kit according to the manufacturer’s instructions, except for skipping the fragmentation step. DNA libraries were pooled and sequenced on a NovaSeq 6000 or NovaSeq X Plus using a 150-cycle kit (paired end, 2 x 150 bp). ChIP-seq analysis was performed similar to what was described previously.^31^ Adapter sequences were trimmed from the read pairs using Cutadapt (-a AGATCGGAAGAGCACACGTCTGAACTCCAGTCA -A AGATCGGAAGAGCGTCGTGTAGGGAAAGAGTGT). Trimmed reads were aligned to the human reference genome (hg38) using BWA (MEM). Properly paired read alignments with mapping scores greater than 30 were used for downstream analysis. PCR duplicates were removed using Picard MarkDuplicates. Peak calling was performed with MACS2 with the options of ‘macs2 callpeak -g hs -f BAMPE --keep-dup all --bdg’. Bigwig files were generated using bedGraphToBigWig tool. Peak metaplots and heatmaps were generated on pooled replicates using deepTools. Gene-specific ChIP enrichment analysis was performed by comparing the number of reads mapped to gene bodies (defined by the featureCounts tool using the GENCODE v42 annotation) in mutant versus WT cell lines. For each biological replicate (individual clone), the read counts of input and pulldown were normalized by total number of aligned reads for the genes whose counts were great than one. Similar to what was previously described,^31^ a generalized linear model was constructed to estimate log2Foldchange of the normalized read counts in pulldown relative to input while adjusting for cell line variability. Shrunken fold change was estimated using the R package DESeq2. The statistical significance of the log2Foldchange was calculated empirically by the R package fdrtool, with the input of Wald statistics calculated using DESeq2. The differential ChIP enrichment between the mutant and WT lines was calculated with an interaction term of cell line (mutant versus WT) and library (pulldown versus input) in a regression model. Plots were generated using ggplot2 and Tableau software. Gene ontology analysis was performed using ShinyGO.

#### RNA isolation, quantification, and quality control

Total RNA was extracted using TRIzol and purified using the Direct-zol RNA MiniPrep kit according to the manufacturer’s instructions. RNA concentration was measured with a Qubit fluorometer (Invitrogen), and RNA quality was determined using a TapeStation (Agilent Technologies).

#### qRT-PCR

Total RNA was reverse-transcribed using the SuperScript IV First-Strand Synthesis System with random hexamers according to the manufacturer’s instructions. RT-qPCR samples were prepared using the SYBR Select Master Mix according to the manufacturer’s instructions. Samples were run in technical duplicates on a QuantStudio 3 Real-Time PCR System (Applied Biosystems). Ct values were normalized against the internal control *GAPDH*. Fold differences in expression levels were calculated according to the 2-ΔΔCt method.^81^

#### RNA-seq

RNA-seq libraries were prepared from 500 ng of total RNA using the KAPA RNA HyperPrep Kit with RiboErase according to the manufacturer’s instructions. Libraries were pooled and sequenced on a NovaSeq 6000 or NovaSeq X Plus using a 150-cycle kit (paired end, 2 x 150 bp).

RNA-seq analysis was performed similar to what was described previously.^31^ Adapter sequences were trimmed from the read pairs using Cutadapt (-a AGATCGGAAGAGCACACGTCTGAACTCCAGTCA -A AGATCGGAAGAGCGTCGTGTAGGGAAAGAGTGT). Trimmed reads were aligned to the human reference genome (hg38) using STAR with GENCODE basic gene annotation (v42). Read counts were calculated as transcripts per million (TPM) using uniquely mapped reads counted using featureCounts in a strand-specific way. Differential expression analysis between the mutant and WT lines was performed by modeling read counts in a generalized linear model accounting for cell lines. Fold change and p-values were calculated using DESeq2, in which two-sided Wald tests for regression coefficients of interest were performed. Differential expression heatmaps were generated using the R package pheatmap. Scatter plots were generated using the Tableau software. Gene ontology analysis was performed using ShinyGO.

#### Recombinant protein expression and purification

Recombinant 4-subunit and 5-subunit PRC2 complexes were expressed and purified similarly to what was previously described.^82^ The 4-subunit PRC2 core complex contains EZH2, SUZ12, EED, and RBBP4 proteins, all of which include a PreScission-protease cleavable N-terminal 6XHis-MBP tags. The 5-subunit AEBP2-PRC2 complex contains an additional PreScission cleavable Strep-II tagged AEBP2 protein. The 5-subunit MTF2-PRC2 and PHF19-PRC2 complexes were reconstituted with the previously generated MTF2 and PHF19 insect cell expression plasmids.^65^ The 6-subunit AEBP2-JARID2-PRC2 complex was made using a single plasmid baculoviral expression system gifted by Dr. Vignesh Kasinath’s lab at University of Colorado Boulder.^16^

Expression vectors based on the pFastBac plasmid system were transformed into DH10Bac competent cells.^13, 31, 65, 82^ Following manufacturer guidelines, baculovirus stocks (P0, P1, and P2) were generated in Sf9 insect cells grown in Sf-900 III SFM medium. P2 titer was determined by a commercially available service based on gp64 expression (Expression Systems). 1 L of Tni insect cells were seeded at 2x10^6^ cells/mL in ESF-921 protein-free medium and infected at a MOI of 1.0 with each baculovirus. Cultures were grown at 27°C and 130 rpm for 66 h before cells were harvested, snap-frozen, and stored at -80°C until purification.

For complex purification, frozen Tni cells were lysed in lysis buffer (10 mM Tris pH 7.5, 250 mM NaCl, 0.5% Igepal, 1mM TCEP) treated with 1X protease inhibitor cocktail. Cell debris was removed by centrifugation at 29,000 for 40 min. Clarified lysate was incubated with amylose resin equilibrated in lysis buffer for 2 h under gentle rotation. Lysate-bead mixture was added to a glass 2.5 cm x 20 cm Econo-Pac chromatography column. The column was washed with 10 column value (cv) of lysis buffer, followed by 16 cv of high salt buffer (10 mM Tris pH 7.5, 500 mM NaCl, 1 mM TCEP), and 16 cv of low salt buffer (10 mM Tris pH 7.5, 150 mM NaCl, 1 mM TCEP). Protein was eluted off the column in 3 cv of elution buffer (low salt buffer, 10 mM maltose). Elution was collected across five fractions. Relevant fractions were combined and concentrated in an Amicon Ultra-15 centrifugal filter unit, 30 kDa MWCO (Millipore Sigma, Cat#UFC903024). The NaCl concentration was increased to ∼250 mM and PreScission protease was added at a mass ratio of 1 part in 50 for overnight cleavage of MBP tags. The protein was then loaded onto a HiTrap Heparin HP 5 x 5 mL affinity column (Cytiva, Cat#17040703) equilibrated in start buffer (10 mM Tris pH 7.5, 150 mM NaCl, 1 mM TCEP). The protein was eluted in a linearly increasing gradient of elution buffer (10 mM Tris pH 7.5, 2 M NaCl, 1 mM TCEP) across 35 cv at a flow rate of 1.5 mL/min. Relevant fractions were combined and concentrated as above, then subsequently loaded onto a HiLoad 16/600 Superdex 200 pg preparative SEC column (Cytiva, Cat#28989335) equilibrated in sizing buffer (20 mM HEPES pH 7.5, 150 mM NaCl, 1 mM TCEP). Protein was eluted in sizing buffer at a flow rate of 0.5 mL/min. Relevant fractions were combined and concentrated as above. The protein was divided into single-use aliquots, which were then snap-frozen and stored at -80 °C until use.

#### Nucleosome reconstitution

Nucleosomes were reconstituted using assembled human histone octamers (Histone Source, Colorado State University, SKU: HOCT_H3C96SC110A_500ug) and a 200-bp 5’-FAM-labeled mononucleosomal DNA containing the 147-bp Widom sequence. Nucleosome reconstitution was performed similarly to what was previously described.^31^ DNA was mixed with each histone octamer in three separate ratios (1:0.8, 1:1, and 1:1.2) in the histone refolding buffer (2 M NaCl, 6 mM Tris pH 7.5, 0.3 mM EDTA and 0.3 mM TCEP). The mixture was first incubated at 37 °C for 30 min, and then the following volumes of reconstitution buffer (20 mM Tris pH 7.5, 1 mM EDTA, 1 mM DTT) were added in 30-min intervals: 10.8 µL, 12 µL, 28 µL, and 64 µL. Nucleosome assembly quality was assessed by gel electrophoresis in a native 6% polyacrylamide 1X TBE gel stained with RedSafe, and the DNA/histone octamer ratio with no unbound DNA was used for downstream binding assays.

#### Methyltransferase activity

Endpoint PRC2 activity was determined using the MTase-Glo Methyltransferase Assay kit. Serial dilutions of PRC2 were prepared in reaction buffer (20 mM Tris pH 8.0, 50 mM NaCl, 1 mM EDTA, 3 mM MgCl2, 0.1 mg/mL BSA, 1 mM DTT) with a reaction concentration starting at 1 µM. S-adenosylmethionine (SAM) and biotinylated mononucleosome were added to each tube to final concentrations of 20 µM and 450 nM, respectively. The reactions were incubated at room temperature for 2 h before being quenched by trifluoroacetic acid to a final concentration of 0.1%. After quenching, MTase-Glo reagent was added to each reaction mixture to a final 1X concentration. After a 30-min incubation, an equal volume of MTase-Glo detection solution was added to each reaction. After another 30-min incubation, luminescence was measured with a TECAN Spark microplate reader. The luminescence of serially diluted S-adenosylhomocysteine (SAH) was run in parallel and a standard curve was generated by linear regression. Net luminescence was determined by subtracting the luminescence from a control containing no protein, and SAH produced by the reaction was determined by using the best-fit equation generated by the SAH standard curve.

#### Fluorescence polarization

Affinities between PRC2 and reconstituted mononucleosomes were determined by fluorescence polarization assays. Various concentrations (4-1000 nM) of the PRC2 complex were prepared by serial dilutions with a final reaction concentration starting at 1 µM in binding buffer (50 mM Tris pH 7.5, 10 mM KCl, 0.1 mM ZnCl2, 2mM BME, 0.1 mg/mL BSA, 5% glycerol). An equal volume of FAM-mononucleosome in the same binding buffer was added to a final concentration of 5 nM per reaction. The binding reactions were incubated at room temperature for 30 min before fluorescence anisotropy was measured with a TECAN Spark microplate reader. The FAM fluorophore was excited at 485 nM with a bandwidth of 20 nM, and emission was measured at 535 nM with a bandwidth of 25 nM. A standard binding curve was generated for each nucleosome with the baseline adjusted to the anisotropy value of the control sample without protein. Apparent *K*_D_^app^ dissociation constant *K*_D_ ^app^ and Hill coefficient were calculated from the mean value of three independent replicates.

#### Cardiomyocyte differentiation of hiPSCs

hiPSCs were differentiated into cardiomyocytes using the GSK3 inhibitor CHIR99021 and Wnt inhibitor IWP2.^30^ Three days before induction of differentiation (day -3), hiPSCs were seeded at 250,000 cells per well of a Geltrex-coated 12-well plate in StemFlex medium with 5 μM ROCK inhibitor Y27632. Medium was replaced daily with StemFlex medium until day 0. On day 0, when cells were around 90% confluent, medium was replaced with RPMI1640 medium containing B-27 supplement (minus insulin) and 10 μM CHIR99021. On day 1, medium was replaced with RPMI1640 medium containing B-27 supplement (minus insulin). On day 3, half of the existing medium was collected from each well and mixed with an equal amount of fresh RPMI1640 medium containing B-27 supplement (minus insulin) and a final concentration of 5 μM IWP2 to replace the remaining media in each well. On day 5, medium was replaced with RPMI1640 medium containing B-27 supplement (minus insulin). On day 7 and every 3 days afterwards, medium was replaced with RPMI1640 medium containing B-27 supplement.

From day 15, cardiomyocytes were further purified using metabolic selection with lactate.^83^ Medium was replaced with lactate selection medium (RPMI1640 medium without glucose, 5 mM sodium DL-lactate, 213 μg/mL L-ascorbic acid, 500 μg/mL human albumin) on day 15 and 17. On day 19, medium was replaced with RPMI1640 medium containing B-27 supplement. On day 22, cells were trypsinized and reseeded at a density of 200,000 cells per well of a Geltrex-coated 12-well plate in RPMI1640 medium containing 10% fetal bovine serum and 5 μM ROCK inhibitor Y27632. From day 23, cardiomyocytes were maintained in RPMI1640 medium containing B-27 supplement with media changes every other day.

Time-lapse images of spontaneous contractions were captured using a Leica DMi8 microscope with a Hamamatsu ORCA-Fusion BT digital camera. Contractions were quantified using the MATLAB-based MotionGUI program.

#### Flow cytometry

Day 8 cardiac progenitor cells and day 15 cardiomyocytes were collected by trypsinization, fixed in 1% formaldehyde in PBS, and washed three times with PBS. Cells were then permeabilized with 90% methanol and washed three times with 0.5% BSA in PBS. Each sample was incubated with anti-cTnT Alexa Fluor 647 or IgG1 κ Alexa Fluor 647 isotype control diluted 1:200 in 0.5% BSA and 0.1% Triton X-100 in PBS for 1 h. Cells were washed twice in 0.5% BSA and 0.1% Triton X-100 in PBS, resuspended in 0.5% BSA in PBS, and filtered through a 70-µm cell strainer. Samples were acquired using a Sony MA900 Cell Sorter. Data were analyzed with FlowJo.

#### Electrophysiological recordings

Spontaneous action potentials in cardiomyocytes were recorded in whole-cell current clamp mode at resting membrane potential. These recordings were made in an extracellular solution (140 mM NaCl, 5.4 mM KCl, 1.8 CaCl2, 1 mM MgCl2, 10 mM glucose, 10 mM HEPES, pH 7.4). Borosilicate patch pipettes with 1.5-2 MΩ resistance were used for recording and filled with internal solution (125 mM K-gluconate, 20 mM KCl, 5 mM NaCl, 1 mM MgCl2, 5 mM MgATP, 10 mM HEPES, pH 7.2). A HEKA EPC 10 amplifier was used for acquiring data and PatchMaster was used to export recorded data. Data were acquired at a sampling rate of 2.4 kHz. Action potential parameters were analyzed using Clampfit.

#### Immunoprecipitation in HEK293T cells

HEK293T cells at around 50% confluency were forward transfected with equal amounts of pcDNA3.1 plasmids expressing either a 3XHA-tagged or a 3XFLAG-tagged protein using Lipofectamine 2000 Transfection Reagent according to the manufacturer’s instructions. Two days after transfection, cells were harvested by trypsinization, washed with cold PBS, and incubated in lysis buffer (25 mM Tris pH 7.5, 150 mM NaCl, 2.5 mM MgCl2, 1% Igepal, 5% glycerol, 2 mM TCEP, 1X protease inhibitor cocktail, 1 µL/mL benzonase) for 30 min on ice. Protein extracts were collected as supernatant after centrifugation at 16,000 x g for 10 min at 4°C. Clarified protein extracts were incubated with ANTI-FLAG M2 Affinity Gel for 2 h at 4°C. Beads were washed five times with wash buffer (25 mM Tris pH 7.5, 150 mM NaCl, 2.5 mM MgCl2, 1% Igepal, 5% glycerol, 2 mM TCEP). Proteins were eluted from the beads in 1X Laemmli sample buffer for 10 min at 95°C and detected by western blotting.

#### Immunoprecipitation in hiPSCs

hiPSCs at around 70-80% confluency were collected by Accutase treatment, washed with cold PBS, and incubated in lysis buffer (25 mM Tris pH 7.5, 150 mM NaCl, 2.5 mM MgCl2, 1% Igepal, 5% glycerol, 2 mM TCEP, 1X protease inhibitor cocktail, 1 µL/mL benzonase) on end-over-end rotation for 30 min at 4°C. Cell lysates were collected as supernatant after centrifugation at 16,000 x g for 10 min at 4°C. Clarified protein extracts were incubated with pre-washed ANTI-FLAG M2 Affinity Gel for 2 h at 4°C. Beads were washed five times with wash buffer (25 mM Tris pH 7.5, 150 mM NaCl, 2.5 mM MgCl2, 1% Igepal, 5% glycerol, 2 mM TCEP). Proteins were eluted by incubating the beads in elution buffer (lysis buffer with 150 ng/µL 3XFLAG peptide) for 30 min at room temperature and collected via centrifugation in Spin-X tubes (Corning, Cat#8161) at 1,000 x g for 2 min at room temperature. The elution was mixed with 4X Laemmli sample buffer for 10 min at 95°C and proteins were detected by western blotting.

### Quantification and statistical analysis

All statistical analyses and definition of replicates are described in the corresponding figure legends. Statistical tests were performed using Microsoft Excel or R. Statistical significance was defined as: * *P* < 0.05, ** *P* < 0.01, and *** *P* < 0.001.

## Acknowledgements

We thank Vigneshwari Kumar, Roxanne Lamanna, and Cyril Gagnon (Weill Cornell) for their contribution at the early stages of the project. We thank Dr. Taeyoung Hwang (Johns Hopkins) for his assistance in our computational analysis. We are grateful for the helpful discussions and feedback from Dr. Jessica Tyler (Weill Cornell) and members of the Long and Pitt labs. We thank Drs. Daniel Youmans and Vignesh Kasinath (University of Colorado Boulder) for kindly providing plasmids. This work is supported by the Weill Cornell Proteomics & Metabolomics Core Facility, the Epigenomics Core Facility and the Genomics Core Facility. YL is supported by R00GM132546 (National Institute of General Medical Sciences), American Heart Association Career Development Award 935361 and Tri-SCI grant 2023-010. GP is supported by R01 HL146149, R01 HL151190, and R01 HL160089 (National Heart, Lung, and Blood Institute).

## Author contributions

Conceptualization, Y.L.; Methodology, C.M. and Y.L.; Investigation, C.M., S.M.R., L.C., J.M.C., Y.L., and A.R.G.; Formal Analysis, Y.L. and C.M.; Writing – Original Draft, Y.L. and C.M.; Visualization, C.M. and Y.L.; Funding Acquisition, Y.L. and G.S.P.; Supervision, Y.L. and G.S.P.

## Supplemental figure legends

**Figure S1.**
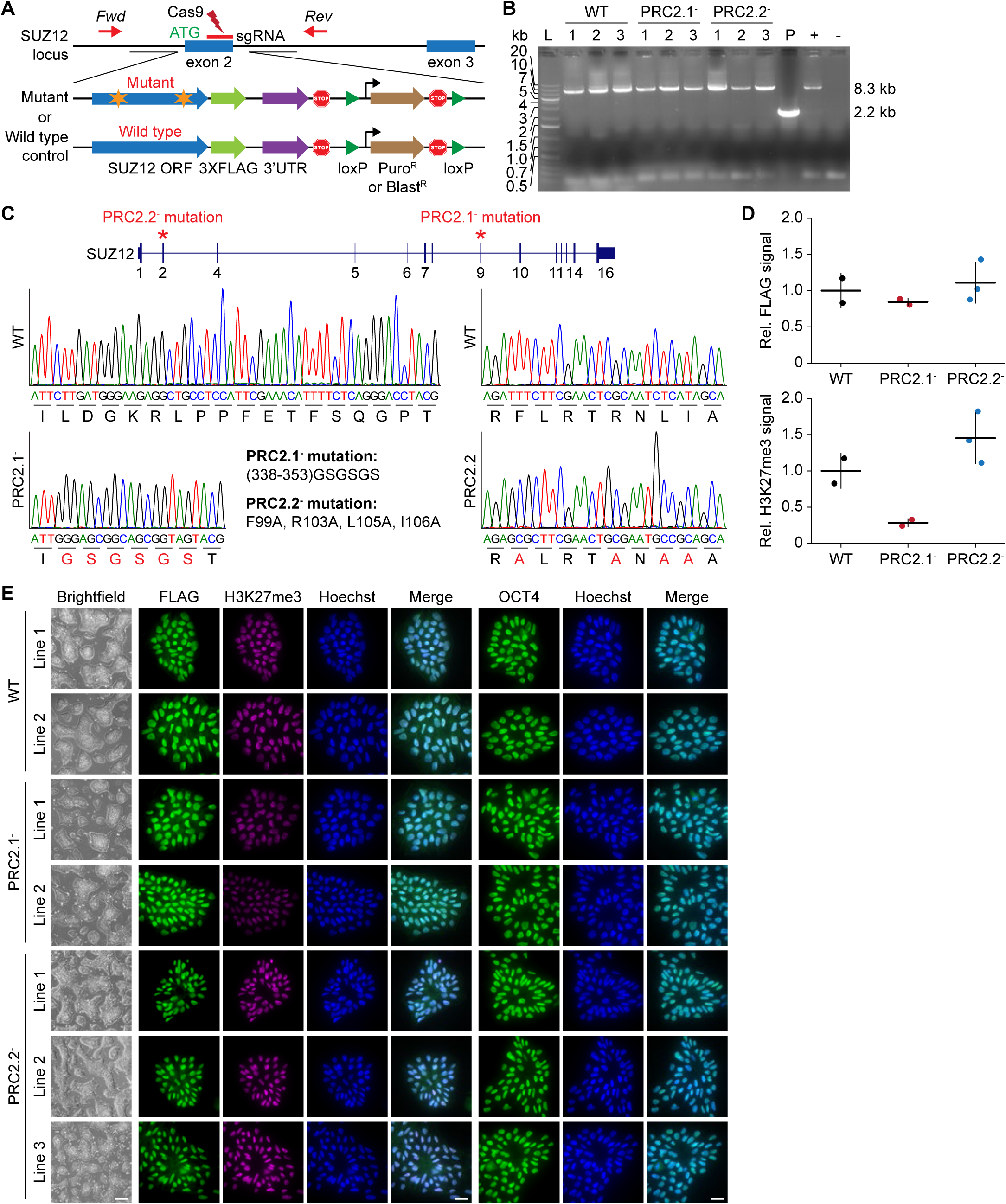
Generation and characterization of PRC2.1 and PRC2.2 separation-of-function mutants, related to Figure 1. (A) Schematic showing the design of WT control and mutant SUZ12 hiPSC cell lines using CRISPR-Cas9 editing. A double-stranded DNA break at the 3’ end of exon 2 of *SUZ12* was generated with sgRNA-Cas9, and the remaining *SUZ12* cDNA along with a C-terminal 3XFLAG epitope tag and *SUZ12* 3’-UTR was inserted immediately downstream of exon 2. To increase the success rate of obtaining homozygous monoclonal lines, two donor plasmids either expressing a puromycin-resistant gene (*Puro^R^*) or a blasticidin S resistant gene (*Blast*^R^) were co-transfected, enabling dual antibiotic selection of cells with both *SUZ12* alleles edited. (B) Genotyping confirming the correct and homozygous modification of the WT and mutant SUZ12 clones. Genomic PCR was performed using the *Fwd* and *Rev* primers indicated in (A). Amplification of the unedited and edited alleles would generate a PCR fragment of 2.2 and 8.3 kb, respectively. Three clones of each edited hiPSC line were analyzed. P represents the parental hiPSC line WTC-11. + represents a positive control clone generated separately. - represents the no template control. L represents the DNA ladder. (C) Sanger sequencing confirming the mutations for the PRC2.1^-^ and PRC2.2^-^ lines using the gel-extracted PCR fragments from (B). (D) Quantification of the FLAG and H3K27me3 signal from the western blot in Figure 1B. The signal was normalized to that of the WT and the β-actin loading control. Data are shown as mean and standard deviation of two (WT and PRC2.1^-^) or three (PRC2.2^-^) biological replicates. (E) Brightfield and immunofluorescence micrographs of WT and mutant SUZ12 hiPSC lines stained with anti-FLAG (green) and anti-H3K27me3 antibody (magenta) as well as anti-OCT4 (green). Nuclei are stained with Hoechst (blue). Scale bar represents 20 µm.

**Figure S2.**
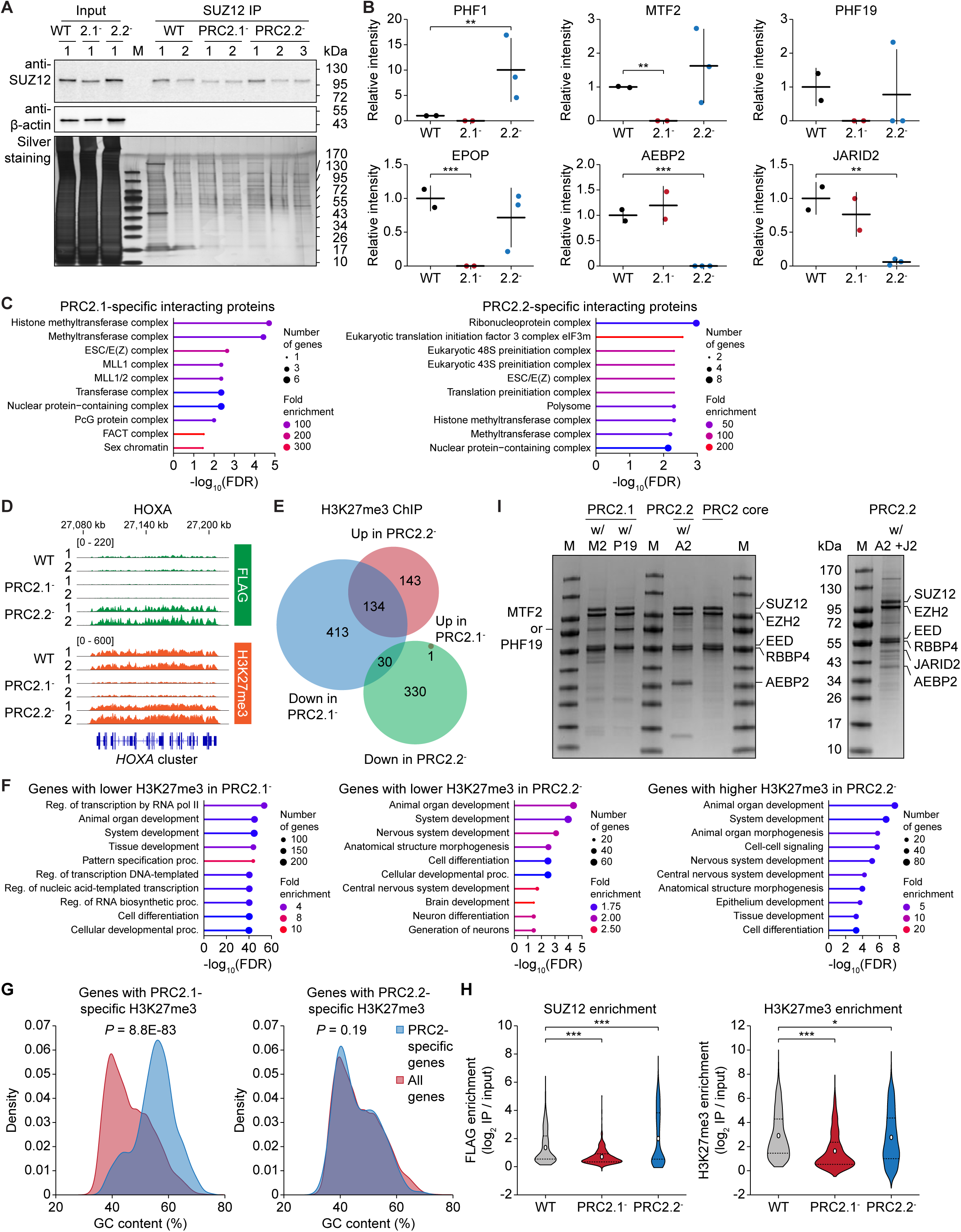
Characterization of PRC2.1 and PRC2.2 subcomplex interactome and target genes, related to Figure 1. (A) Western blot and silver staining of the SUZ12 immunoprecipitated samples in WT and mutant SUZ12 hiPSC lines. Whole cell lysate (input) was incubated with anti-SUZ12 beads to pull down SUZ12. Two or three homozygous clones from each edited line were used. M represents the protein ladder. (B) Relative mass spectrometry intensity of co-immunoprecipitated PRC2 accessory proteins. Horizonal bar represents the mean and vertical bar represents the standard deviation. Data are shown as mean and standard deviation of two (WT and PRC2.1^-^) or three (PRC2.2^-^) biological replicates. ** *P* < 0.01, *** *P* < 0.001, two-tailed Student’s *t*-test. (C) Gene ontology analysis of SUZ12-interacting proteins that are specifically found in PRC2.1 or PRC2.2. (D) FLAG and H3K27me3 ChIP-seq genome tracks of the WT and mutant SUZ12 hiPSC lines at the *HOXA* cluster. (E) Venn diagram showing the number of genes with increasing or decreasing H3K27me3 levels upon loss of PRC2.1 and PRC2.2 (all genes are Sig or SigFC in Figure 1F, and significantly enriched between IP and input in the WT line). This is an extended version of Figure 1G, with the addition of genes that gain H3K27me3 upon loss of PRC2.1 or PRC2.2. (F) Gene ontology analysis of PRC2.1-repressed genes (blue cycle in (E)), PRC2.2-repressed genes (green circle in (E)), and PRC2.2-activated genes (red circle in (E)). (G) Histograms showing the GC content of genes with PRC2.1- (left) and PRC2.2-specific (right) H3K27me3 deposition as compared to that of all genes. (H) Violin plots showing the enrichment of FLAG-tagged SUZ12 (left) and H3K27me3 (right) enrichment in WT and mutant SUZ12 hiPSCs. This is an alternative representation of the FLAG and H3K27me3 ChIP-seq results in Figure 1E. * *P* < 0.05, *** *P* < 0.001, Welch’s *t*-test. (I) SDS-PAGE gel showing the components of recombinant PRC2 subcomplexes. M2: MTF2, P19: PHF19, A2: AEBP2, J2: JARID2. Short isoforms of AEBP2 (amino acid 201-503) and JARID2 (amino acid 118-450) were used. M represents the protein ladder.

**Figure S3.**
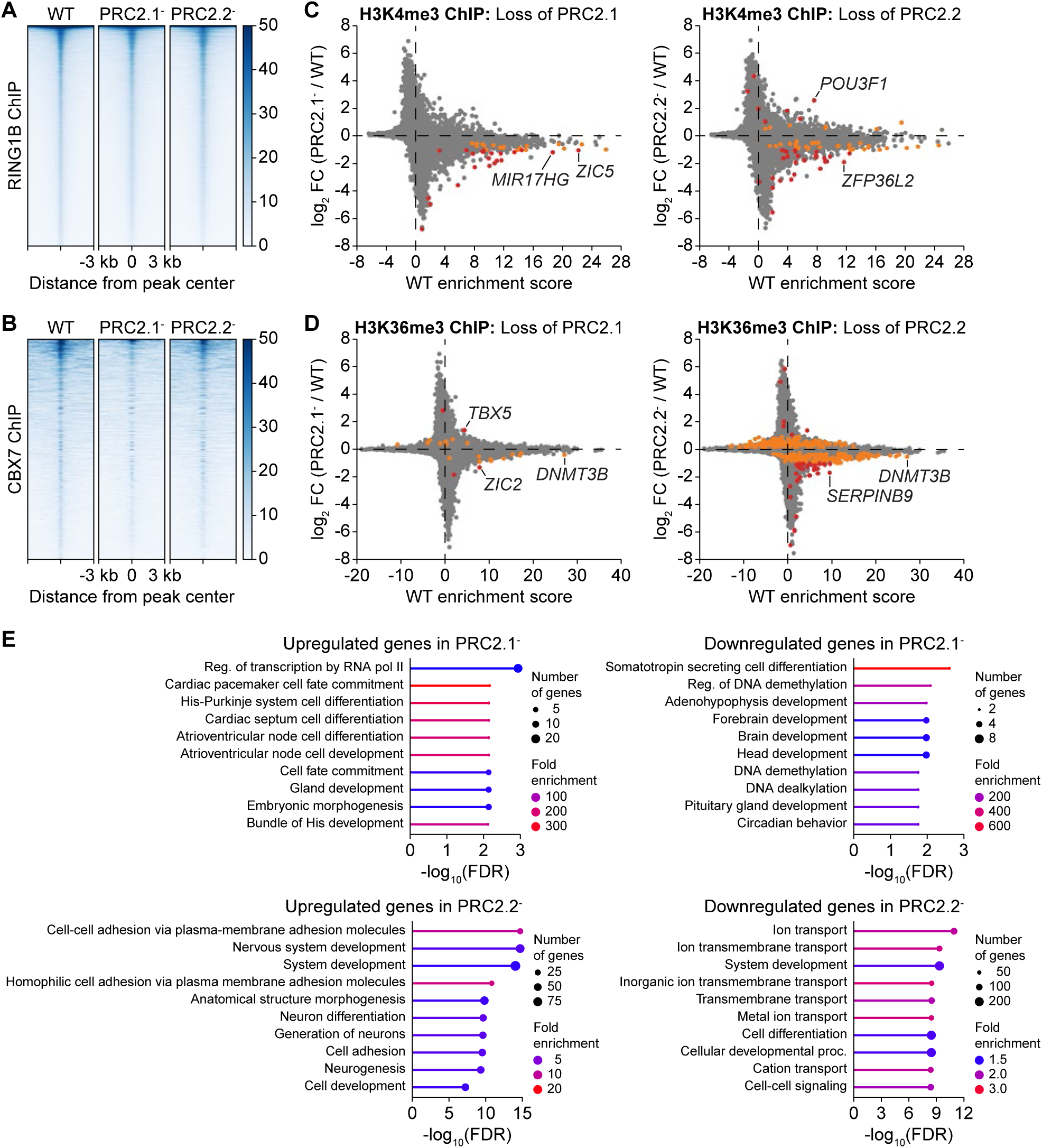
Epigenomic and transcriptomic changes upon loss of PRC2 subcomplexes, related to. Figure 2 (**A-B**) ChIP-seq heatmaps showing RING1B (A) and CBX7 (B) chromatin occupancy upon loss of PRC2.1 or PRC2.2. All regions in the heatmaps are sorted by decreasing peak signal based on the WT peak profile. (**C-D**) ChIP-seq gene scatter plots comparing genome-wide chromatin occupancy changes of H3K4me3 (C) and H3K36me3 (D) upon loss of PRC2.1 or PRC2.2. Two independent clones of each genotype (n = 2) were compared by empirical Wald tests for individual genes. Grey dots (Insig) indicate a multiple test-corrected (FDR) *p*-value ≥ 0.1, orange dots (Sig) an FDR-adjusted *p*-value < 0.1 and |log2FC| < 1, and red dots (SigFC) an FDR-adjusted *p*-value < 0.1 and |log2FC| ≥ 1. (**E**) Gene ontology analysis of up- and downregulated genes (sigFC genes in RNA-seq of Figure 2F) upon loss of PRC2.1 or PRC2.2.

**Figure S4.**
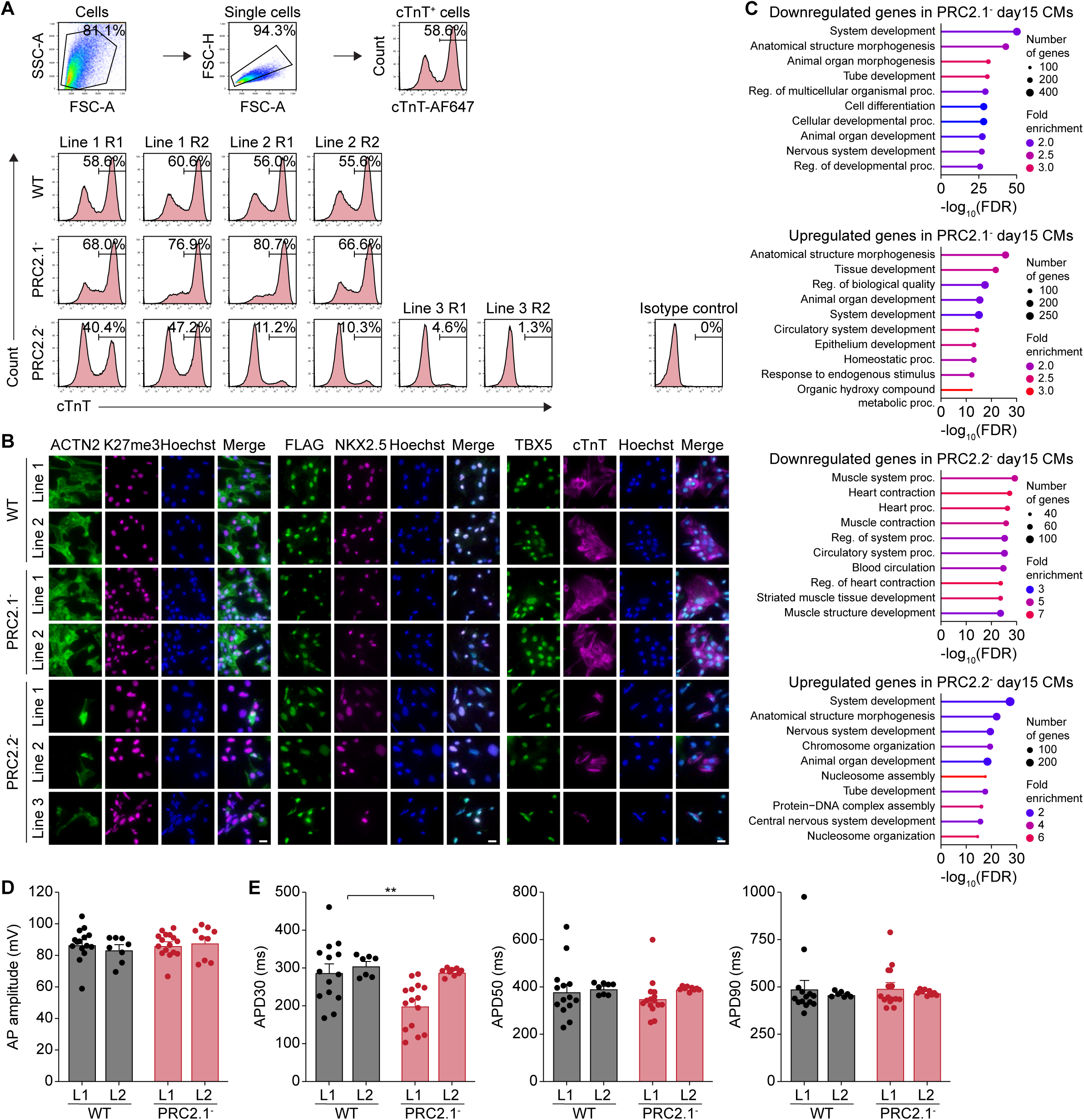
Characterization of cardiomyocytes upon loss of PRC2.1 and PRC2.2, related to Figure 3. (**A**) Flow cytometry histograms showing cTnT expression of WT and mutant SUZ12 day 15 cardiomyocytes. The gating logic is depicted on the left. Two independent replicates of each line are shown. (**B**) Immunofluorescence micrographs of WT and mutant SUZ12 day 15 cardiomyocytes stained with anti-ACTN2 (green) and anti-H3K27me3 antibody (magenta), anti-FLAG (green) and anti-NKX2-5 (magenta), as well as anti-TBX5 (green) and anti-cTnT (magenta). Nuclei are stained with Hoechst (blue). Scale bar represents 20 µm. (**C**) Gene ontology analysis of up- and downregulated genes (sigFC genes in RNA-seq of Figure 3G) upon loss of PRC2.1 or PRC2.2 in day 15 cardiomyocytes. (**D-E**) Bar plots showing action potential amplitude (D) the action potential duration APD30, ADP50, and APD90 (E) in WT and PRC2.1^-^ day 24 cardiomyocytes. Data are shown as mean and standard deviation of 15 (WT replicate 1), 8 (WT replicate 2), 15 (PRC2.1^-^ replicate 1), and 9 (PRC2.1^-^ replicate 2) cell measurements. ** *P* < 0.01, two-way ANOVA.

**Figure S5.**
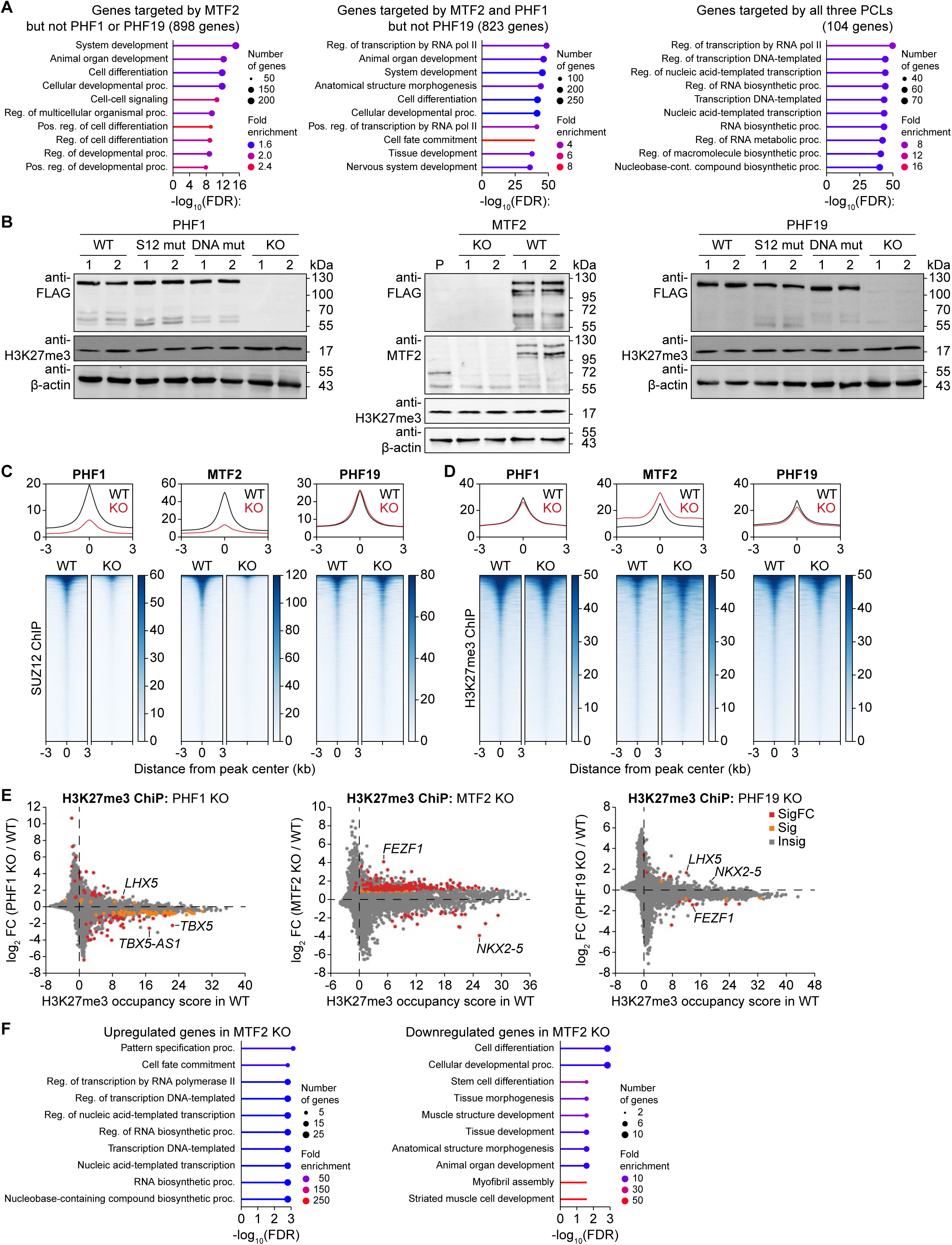
Epigenomic changes upon PCL protein depletion, related to Figure 4. (**A**) Gene ontology analysis of PCL protein target genes from the Venn diagram of Figure 4D. (**B**) Western blot on whole-cell lysates of WT, SUZ12- and DNA-binding mutant, and KO PHF1 (left), MTF2 (middle) and PHF19 (right) hiPSC lines. Note that the shift in molecular weight between the parental (P) and WT line of MTF2 is caused by the addition of the 3XFLAG-HALO tag. (**C-D**) ChIP-seq metaplots (top) and heatmaps (bottom) showing SUZ12 chromatin occupancy (C) and H3K27me3 levels (D) upon depletion of each PCL protein. All regions in the heatmaps are sorted by decreasing peak signal based on the WT peak profile. (**E**) ChIP-seq gene scatter plots comparing genome-wide changes of H3K27me3 levels upon loss of PHF1, MTF2, or PHF19. Two independent clones of each genotype (n = 2) were compared by empirical Wald tests for individual genes. Grey dots (Insig) indicate a multiple test-corrected (FDR) *p*-value ≥ 0.1, orange dots (Sig) an FDR-adjusted *p*-value < 0.1 and |log2FC| < 1, and red dots (SigFC) an FDR-adjusted *p*-value < 0.1 and |log2FC| ≥ 1. (**F**) Gene ontology analysis of the up- and downregulated genes in MTF2 KO hiPSCs.

**Figure S6.**
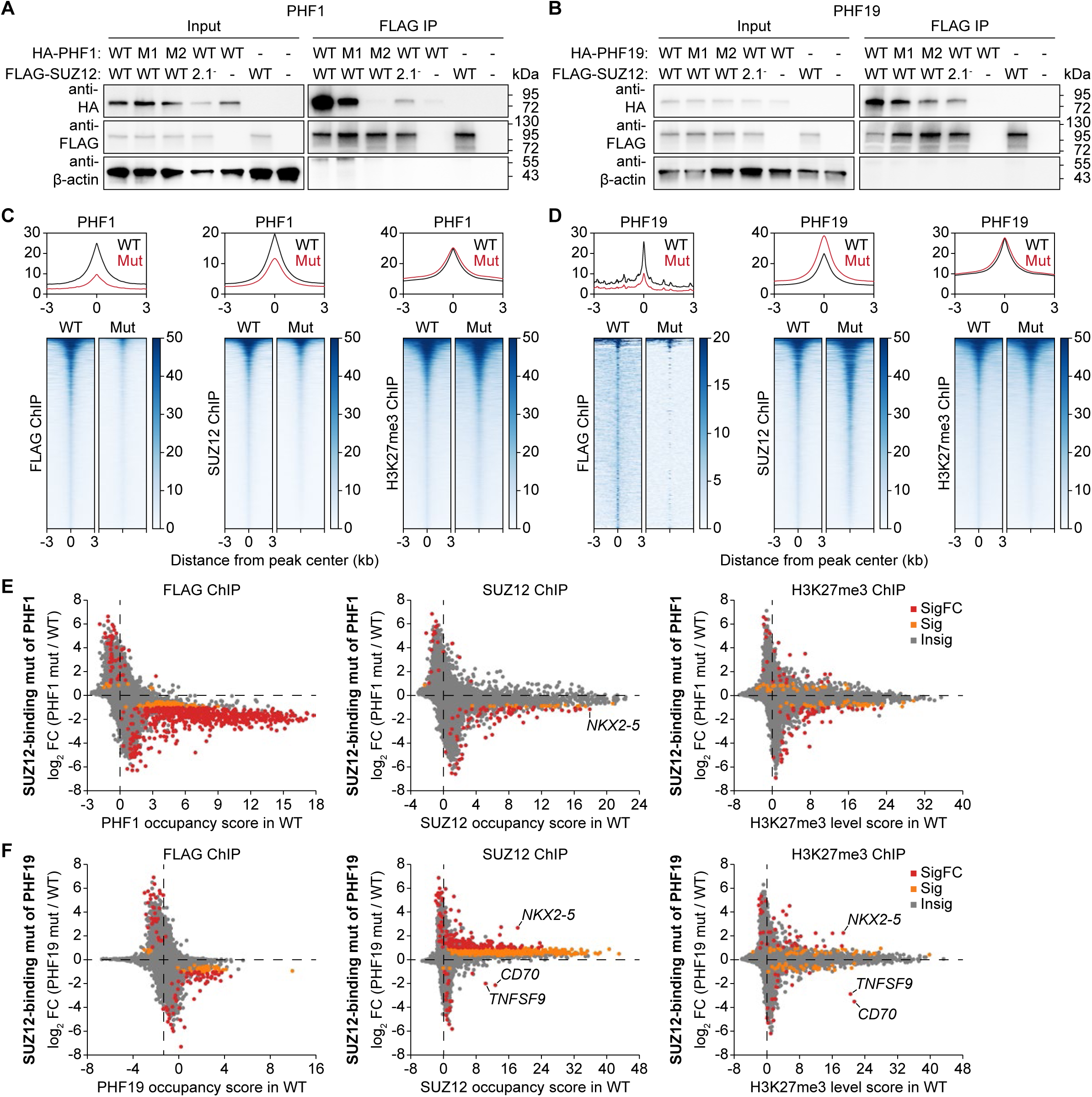
Characterization of the PHF1 and PHF19 SUZ12-binding mutants, related to Figure 5. (**A-B**) Western blot of co-immunoprecipitation assay performed in HEK293T cells by transiently expressing ectopic HA-tagged PHF1 (A) or PHF19 (B) and FLAG-tagged SUZ12. Input samples represent the whole cell lysate and the FLAG-IP samples the fraction eluted off the anti-FLAG beads. Mutants of PHF1: M1: L558A, W561A; M2: R548A, L558A, W561A. Mutants of PHF19: M1: L571A, W574A; M2: R561A, L571A, W574A. PRC2.1^-^ mutant of SUZ12: (338-353)GSGSGS. (**C-D**) ChIP-seq metaplots (top) and heatmaps (bottom) showing FLAG, SUZ12 and H3K27me3 chromatin occupancy for WT and SUZ12-binding mutant PHF1 (C) and PHF19 (D) hiPSCs lines. All regions in the heatmaps are sorted by decreasing peak signal based on the WT peak profile. (**E-F**) ChIP-seq gene scatter plots comparing genome-wide chromatin occupancy changes of FLAG-tagged PHF1/19, SUZ12, and H3K27me3 between WT and SUZ12-binding mutant PHF1 (E) and PHF19 (F). Two independent clones of each genotype (n = 2) were compared by empirical Wald tests for individual genes. Grey dots (Insig) indicate a multiple test-corrected (FDR) *p*-value ≥ 0.1, orange dots (Sig) an FDR-adjusted *p*-value < 0.1 and |log2FC| < 1, and red dots (SigFC) an FDR-adjusted *p*-value < 0.1 and |log2FC| ≥ 1.

**Figure S7.**
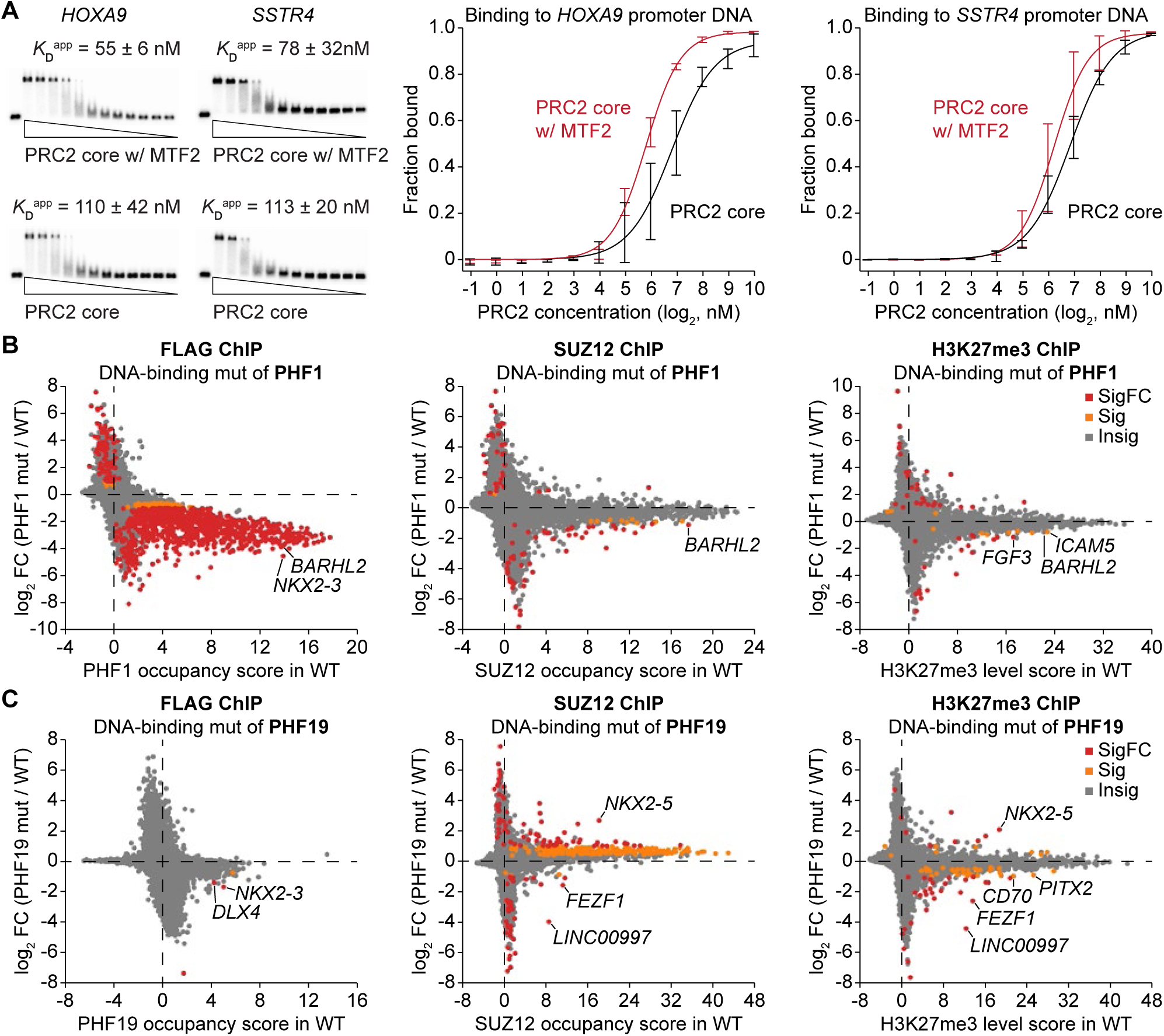
Characterization of the PCL protein DNA-binding mutants, related to Figure 6. (**A**) EMSA to compare the binding affinity between an *HOXA9* and *SSTR4* promoter DNA and the recombinant PRC2 core complex or the MTF2-containing PRC2.1 complex. Representative EMSA gels are shown on the left. Quantification of the fraction of DNA bound with error bars representing the standard deviation from three independent experiments is shown on the right. (**B-C**) ChIP-seq gene scatter plots comparing genome-wide chromatin occupancy changes of FLAG-tagged PHF1/19, SUZ12, and H3K27me3 between WT and DNA-binding mutant PHF1 (B) and PHF19 (C). Two independent clones of each genotype (n = 2) were compared by empirical Wald tests for individual genes. Grey dots (Insig) indicate a multiple test-corrected (FDR) *p*-value ≥ 0.1, orange dots (Sig) an FDR-adjusted *p*-value < 0.1 and |log2FC| < 1, and red dots (SigFC) an FDR-adjusted *p*-value < 0.1 and |log2FC| ≥ 1.

**Figure S8.**
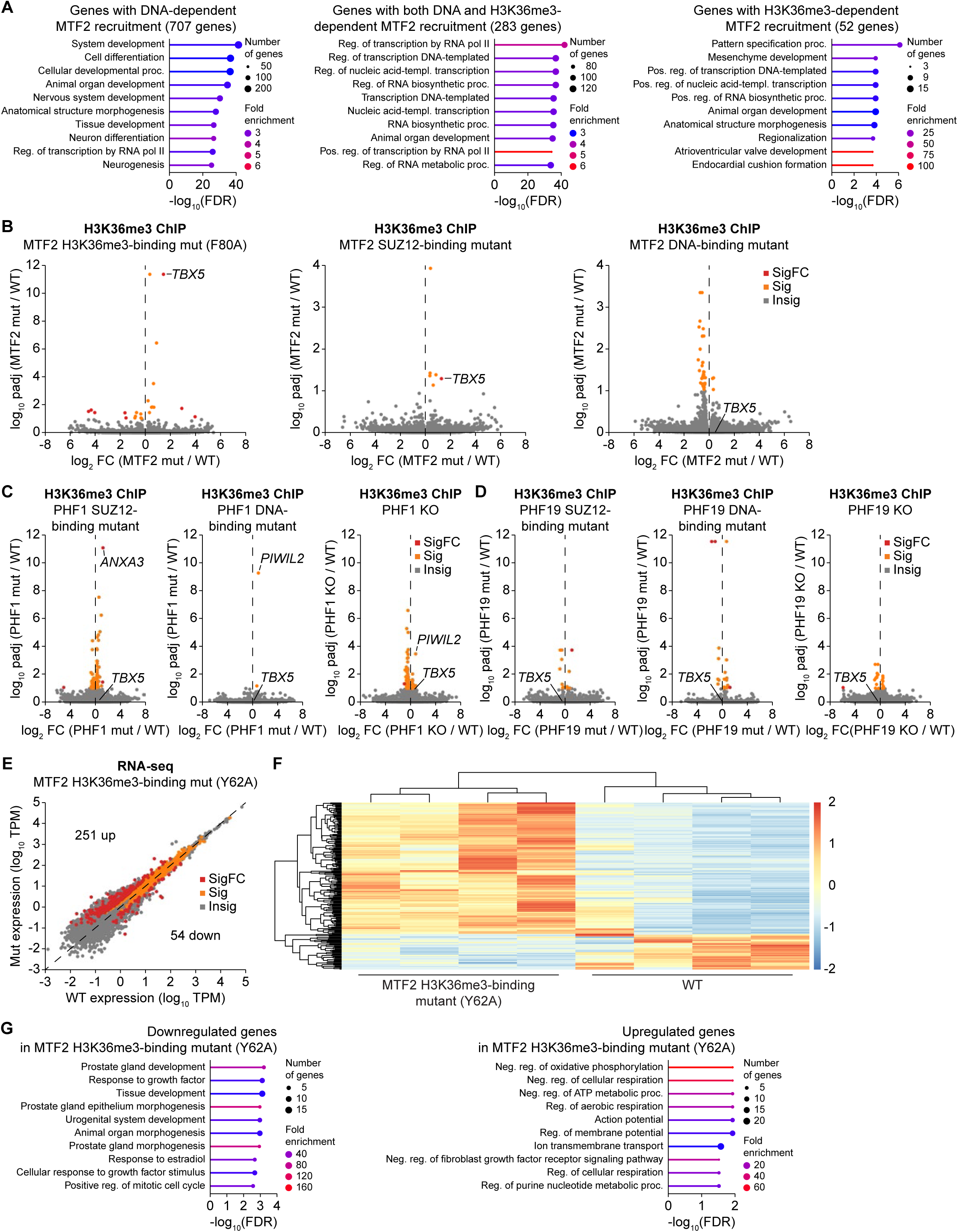
Characterization of the MTF2 H3K36me3-binding mutant, related to Figure 7. (**A**) Gene ontology analysis of the genes with DNA- and H3K36me3-specific recruitment of MTF2 as illustrated in the Venn diagram in Figure 7C (left, FLAG-MTF2 ChIP-seq). (**B-D**) ChIP-seq volcano plots comparing the gene-specific changes of H3K36me3 levels between WT and a second H3K36me3-binding mutant of MTF2 (F80A, B, left), the SUZ12-binding mutant of MTF2 (B, middle), the DNA-binding mutant of MTF2 (B, right), the SUZ12-binding mutant of PHF1 (C, left), the DNA-binding mutant of PHF1 (C, middle), the PHF1 KO (C, right), the SUZ12- binding mutant of PHF19 (D, left), the DNA-binding mutant of PHF19 (D, middle), and the PHF19 KO (D, right). Grey dots (Insig) indicate a multiple test-corrected (FDR) *p*-value ≥ 0.1, orange dots (Sig) an FDR-adjusted *p*-value < 0.1 and |log2FC| < 1, and red dots (SigFC) an FDR-adjusted *p*-value < 0.1 and |log2FC| ≥ 1. (E) RNA-seq scatter plots showing gene expression changes between WT and H3K36me3- binding mutant (Y62A) MTF2 hiPSCs. Two independent clones of each genotype cultured independently twice (n = 4) were compared for statistical significance (two-sided Wald test). Grey dots (Insig) indicate a multiple test-corrected (FDR) *p*-value ≥ 0.05, orange dots (Sig) an FDR-adjusted *p*-value < 0.05 and |log2FC| < 1, and red dots (SigFC) an FDR-adjusted *p*-value < 0.05 and |log2FC| ≥ 1. Number of up- and downregulated genes (SigFC, red dots) are indicated. (F) Gene cluster heatmap showing the gene expression changes between WT and H3K36me3- binding mutant (Y62A) MTF2 hiPSC lines. All depicted genes are SigFC genes (red dots in (E)). (G) Gene ontology analysis of down- or upregulated genes in the RNA-seq (SigFC genes, red dots in (E)).

**Figure S9.**
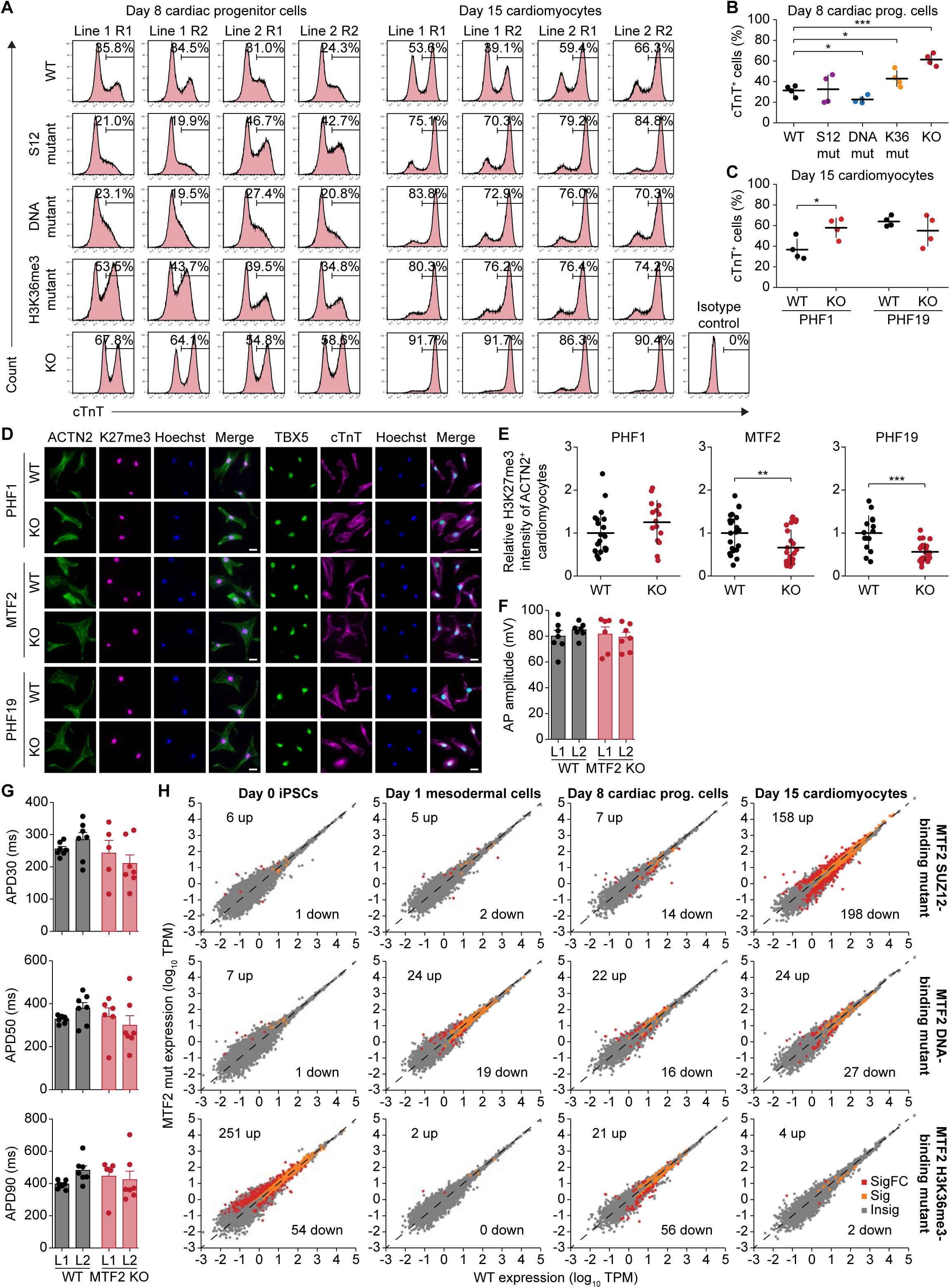
Characterization of PCL protein variant cardiomyocytes, related to Figure 8. (A) Flow cytometry histograms showing cTnT expression of WT and variant MTF2 day 8 cardiac progenitor cells and day 15 cardiomyocytes. Two independent replicates of each line are shown. (B) Percentage of WT and variant MTF2 cells expressing cTnT after 8 days of differentiation as determined by flow cytometry. Data are shown as mean and standard deviation of four biological replicates for each genotype. * *P* < 0.05, *** *P* < 0.001, two-tailed Student’s *t*-test. (C) Percentage of PHF1 and PHF19 WT and KO cells expressing cTnT after 15 days of differentiation as determined by flow cytometry. Data are shown as mean and standard deviation of four biological replicates for each genotype. * *P* < 0.05, two-tailed Student’s *t*-test. (D) Immunofluorescence micrographs of PHF1, MTF2, and PHF19 WT and KO day 25 cardiomyocytes stained with anti-ACTN2 (green) and anti-H3K27me3 antibody (magenta) as well as anti-TBX5 (green) and anti-cTnT (magenta). Nuclei are stained with Hoechst (blue). Scale bar represents 20 µm. (E) Quantification of the H3K27me3 signal of ACTN2^+^ day 15 cardiomyocytes from the immunofluorescence micrographs in (D). Data are shown as mean and standard deviation of 20 (PHF1 WT), 15 (PHF1 KO), 23 (MTF2 WT), 25 (MTF2 KO), 15 (PHF19 WT), and 26 (PHF19 KO) quantified cells. ** *P* < 0.01, *** *P* < 0.001, two-tailed Student’s *t*-test. (**F-G**) Bar plots showing action potential amplitude (F) and the action potential duration APD30, ADP50, and APD90 (G) in WT and MTF2 KO day 24 cardiomyocytes. Data are shown as mean and standard deviation of seven cell measurements. (H) RNA-seq scatter plots showing gene expression changes between WT and mutant MTF2 lines at day 0, 1, 8, and 15 of the cardiac differentiation process. Two independent clones of each genotype cultured independently twice (n = 4) were compared for statistical significance (two-sided Wald test). Grey dots (Insig) indicate a multiple test-corrected (FDR) *p*-value ≥ 0.05, orange dots (Sig) an FDR-adjusted *p*-value < 0.05 and |log2FC| < 1, and red dots (SigFC) an FDR-adjusted *p*-value < 0.05 and |log2FC| ≥ 1. Number of up- and downregulated genes (SigFC, red dots) are indicated.

